# Cellular selectivity of STING stimulation determines priming of anti-tumor T cell responses

**DOI:** 10.1101/2021.12.01.469893

**Authors:** Bakhos Jneid, Aurore Bochnakian, Fabien Delisle, Emeline Djacoto, Jordan Denizeau, Christine Sedlik, Frédéric Fiore, Robert Kramer, Ian Walters, Sylvain Carlioz, Bernard Malissen, Eliane Piaggio, Nicolas Manel

## Abstract

T cells that recognize tumor antigens are crucial for anti-tumor immune responses. Induction of anti-tumor T cells in immunogenic tumors depends on STING, the intracellular innate immune receptor for cyclic guanosine monophosphate-adenosine monophosphate (cGAMP) and related cyclic dinucleotides (CDNs). However, the optimal way to leverage STING activation in non-immunogenic tumors is still unclear. Here, we show that cGAMP delivery by intra-tumoral injection of virus-like particles (cGAMP-VLP) leads to differentiation of tumor-specific T cells, decrease in tumor regulatory T cells (Tregs) and anti-tumoral responses that synergize with PD1 blockade. By contrast, intra-tumoral injection of synthetic CDN leads to tumor necrosis and systemic T cell activation but no differentiation of tumor-specific T cells, and a demise of immune cells in injected tumors. Analyses of cytokine responses and genetic models revealed that cGAMP-VLP preferentially targets STING in dendritic cells at a 1000-fold less dose than synthetic CDN. Sub-cutaneous administration of cGAMP-VLP showed synergy when combined with a tumor Treg-depleting antibody to elicit systemic tumor-specific T cells, leading to complete and lasting tumor eradication. These finding show that cell targeting of STING stimulation shapes the anti-tumor T cell response and reveal a therapeutic strategy with T cell modulators.

## Introduction

T cells that recognize tumor antigens are critical effectors of the anti-tumor immune response. Most cancer patients do not naturally mount effective T cell responses against their tumors. Immune-checkpoint blocking antibodies (ICB) led to remarkable therapeutic success albeit in a fraction of patients and tumor types. ICB require pre-existing anti-tumor T cells responses to work (Tumeh et al., 2014). The understanding of the mechanisms that efficiently generate anti-tumor T cells has the potential to expand the efficacy of ICB by enabling new classes of immunotherapeutic agents.

Specialized antigen presenting-cells can stimulate T cell responses from naive cells. Antigen-presenting cells are activated by innate immune signals emanating from germline-encoded pattern recognition receptors that recognize non-self or altered-self molecules. STING is an intracellular pattern recognition receptor for cyclic dinucleotides (CDNs) implicated in the response to bacteria and to intracellular DNA of foreign and altered-self origins. In mouse models, spontaneous generation of anti-tumor T cells against immunogenic tumors has been shown to rely on STING activation (Woo et al., 2014). Intra-tumoral injection of synthetic CDNs that activate STING stimulate anti-tumor responses, but the underlying mechanisms remain unclear (Corrales et al., 2015). In fact, synthetic CDNs can have contradictory immune-stimulatory and immuno-ablative effects at different doses (Sivick et al., 2018). Given that STING is broadly expressed in normal tissues and also tumors, the potential for tissue-specific activation of STING may either support protective or pathological responses (Liu et al., 2014). For example, STING activation within T cells inhibits their proliferation and, at least in mouse, triggers their death by apoptosis (Cerboni et al., 2017; Gulen et al., 2017). The optimal cell type for STING activation with the aim of priming antigen-specific anti-tumor T cell responses is unknown.

The STING pathway also plays an evolutionary conserved role in anti-viral immunity (Goto et al., 2020; Morehouse et al., 2020). Moreover, the natural mammalian STING agonist, 2’3’-cGAMP (cGAMP) can be packaged in particles of enveloped viruses, leading to STING activation in target cells immediately after fusion of the viral particles (Bridgeman et al., 2015; Gentili et al., 2015). This represents a Trojan horse system of antiviral defense without the need to detect viral nucleic acids. Consequently, cGAMP can be packaged in non-infectious enveloped virus-like particles (VLP). These enveloped retroviral VLPs can be readily produced and purified, enabling the production of cGAMP-containing VLPs (cGAMP-VLP) (Bridgeman et al., 2015; Gentili et al., 2015). Inclusion of cGAMP enhances the immunogenicity of VLPs displaying influenza virus or SARS-CoV-2 glycoproteins (Chauveau et al., 2021).

Here, we leveraged the biological properties of cGAMP-VLP to investigate anti-tumoral immunity induced by STING activation. We characterized STING activation *in vivo* by cGAMP-VLP compared to established synthetic cyclic dinucleotide (CDN). Using cGAMP-VLP, we show that STING is essential in dendritic cells for the induction of tumor-specific T cell responses that respond to ICB. Finally, we identify a critical role of tumor Treg in limiting anti-tumor T cell response induced by STING activation.

## Results

### Production and characterization of cGAMP-VLP

cGAMP-VLP were produced by transient transfection of 293FT cells and purified through a sucrose cushion and two rounds of ultra-centrifugation. We routinely measured the concentration of cGAMP and of p24 (antigen of the structural viral protein Gag of HIV-1 used to produce the VLP) in the purified preparations. Using a nanoparticle tracker, we observed a homogenous distribution average at 158 nm, which is consistent with the size of retroviral particles (**Figure S1A**). We visualized the cGAMP-VLP by electron microscopy, which confirmed the size range (**Figure S1B**). Titration of the cGAMP-VLP on THP-1 cells induced a dose-dependent upregulation of SIGLEC-1, an IFN-stimulated gene that is upregulated in response to STING activation (**Figure S1C**). Comparison to the clinically tested CDN ADU-S100 (Corrales et al., 2015) or to synthetic 2’3’-cGAMP demonstrated that cGAMP-VLP was ∼500x and ∼200x more effective, respectively. We enhanced intracellular delivery of ADU-S100 or 2’3’-cGAMP using lipofectamine. cGAMP-VLP was still ∼9x and ∼50x more effective than the lipofected ADU-S100 or 2’3’-cGAMP, respectively.

### Intra-tumoral injection of cGAMP-VLP induces tumor rejection

To assess the anti-tumor effect of cGAMP-VLP, we used the male murine tumor MB49 which can be rejected by T cell responses (Perez-Diez et al., 2007). We initiated treatment on 50 mm^3^ tumors and performed three intra-tumoral injections of cGAMP-VLP containing 50 ng cGAMP or injections of PBS, every three days (**Figure S1D**). Tumors grew continuously in the PBS group, and a minority of mice (3/8) spontaneously eliminated the tumor (**Figure S1E**). In contrast, all mice treated with cGAMP-VLP (8/8) eradicated the tumor. cGAMP-VLP induced a statistically significant anti-tumor effect (**Figure S1F**). We also measured the tumor-specific T cell response in the blood in some mice. cGAMP-VLP induced a significant increase in the CD4+ T cells responding to the tumor antigen DBy (**Figure S1G**). In addition, a fraction of mice treated with cGAMP-VLP showed a high level of CD8^+^ T cell responses to the tumor antigen Uty.

### Intra-tumoral injection of cGAMP-VLP induces T cell responses in a poorly immunogenic tumor model

This result suggested that cGAMP-VLP has the capacity to stimulate T cell responses against tumor antigens. To investigate this effect, we switched to the murine tumor B16-OVA, which is poorly responsive to PD1 blockade (De Henau et al., 2016). We started treatment on palpable tumors and performed three intra-tumoral injections of either cGAMP-VLP, empty VLP (VLP), empty VLP with the matched dose of free 2’3’-cGAMP co-injected (VLP + equivalent cGAMP), free 2’3’-cGAMP alone, free ADU-S100 or PBS (**Figure 1A**). For cGAMP-VLP, we used an injection dose containing 33 ng of cGAMP in one experiment and 50 ng in a second experiment. For free 2’3’-cGAMP and ADU-S100, we used 50 µg per injection. To evaluate STING activation, we measured cytokines in the serum 3h after the first injection (**Figure 1B**). cGAMP-VLP, ADU-S100 and 2’3’-cGAMP induced IFN-α, IFN-β, IL-6 and TNF-α. Empty VLP did not induce these cytokines. cGAMP-VLP induced significantly more IFN-α, IFN-β and TNF-α than the VLP + equivalent cGAMP, consistent with the enhanced intra-cellular delivery of cGAMP contained in the VLP of cGAMP-VLP. Low (33 ng) or higher doses (50 ng) cGAMP-VLP induced similar levels of cytokines compared to 50 µg free 2’3’-cGAMP. ADU-S100 (50 µg) induced higher levels of the cytokine, suggesting that STING stimulation across cell types was not saturated by cGAMP-VLP. These results show that cGAMP-VLP induces cytokine responses that require a 1000-fold less amount of cGAMP compared to the synthetic molecule.

**Figure 1.**
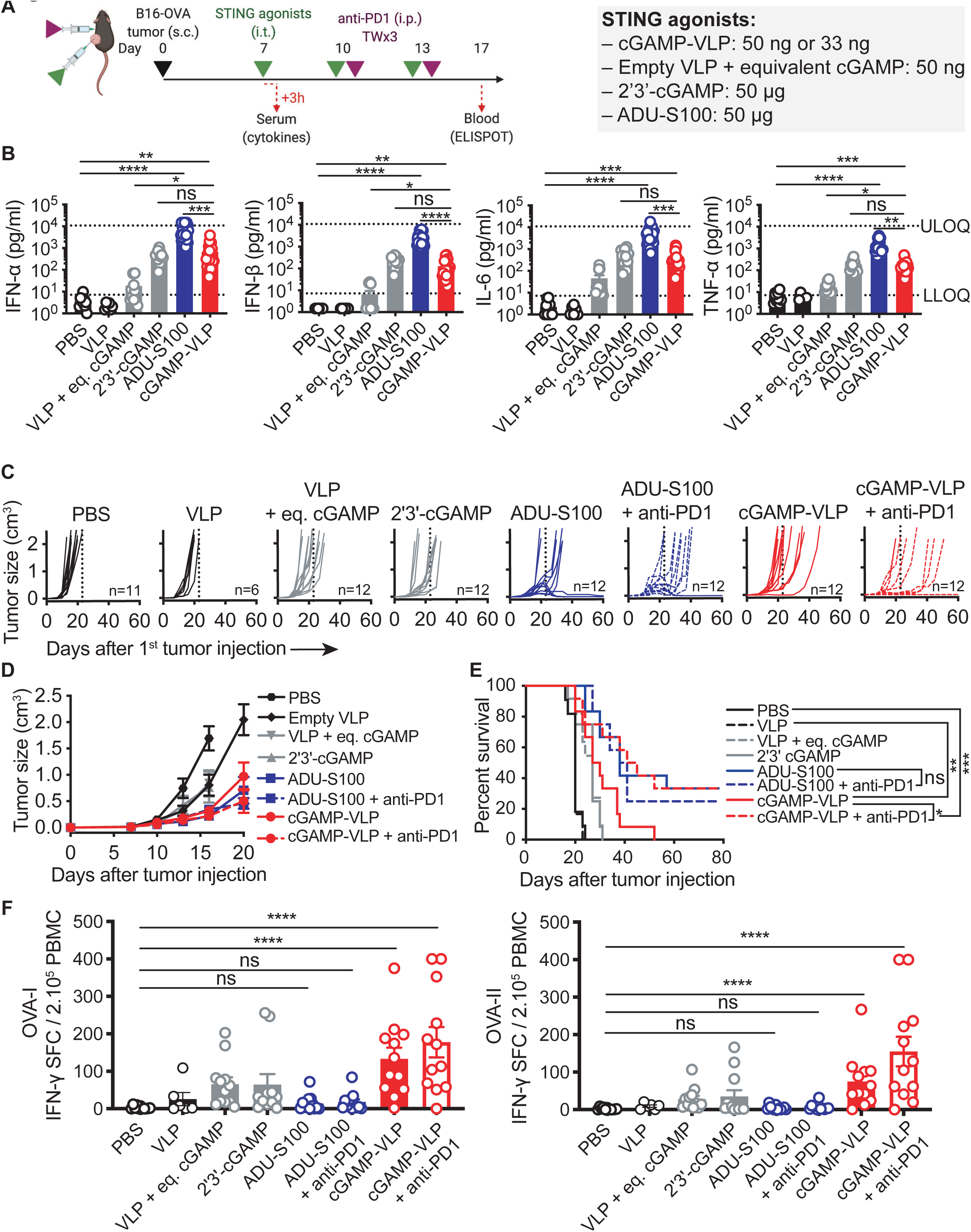
cGAMP-VLP induces tumor-specific T cell responses in a non-immunogenic tumor model. **(A)** Overview of the experimental design (TW = twice weekly). Treatments were initiated on palpable tumors (15-20 mm^3^ range). **(B)** Concentrations of IFN-α, IFN-β, IL-6 and TNF-α in the serum of B16-OVA tumor-bearing mice 3 hours after treatment (bar at mean + SEM, n = 6 to 24 mice per group, combined from 2 independent experiments, Kruskal-Wallis with Dunn post-test, LLOQ = lower limit of quantification, ULOQ = upper limit of quantification). **(C)** Growth curves of individual B16-OVA tumors treated as indicated. Vertical dotted line indicates the death of the last mouse in the PBS-injected group. **(D)** Mean growth over time of B16-OVA tumors treated as indicated (line at mean + SEM, n = 6 to 12 mice per group, combined from 2 independent experiments). **(E)** Survival of B16-OVA tumor-bearing mice treated as indicated (log-rank Mantel-Cox test). **(F)** Ova-specific CD8 (OVA-I) and CD4 (OVA-II) T cell responses in blood, assessed by IFN-γ ELISPOT (bar at mean + SEM, n = 6 to 12 mice per group, combined from 2 independent experiments, Kruskal-Wallis test with Dunn post-test).

We next measured tumor growth. ADU-S100 and cGAMP-VLP were tested with or without anti-PD1 to assess the impact of immune checkpoint inhibition on the response. cGAMP-VLP induced a delay in tumor growth (**Figure 1C, 1D**). Adding anti-PD1 enhanced this delay and led to complete responses in a subset of mice (**Figure 1C**). In comparison, ADU-S100 induced a delay in tumor progression and some complete responses, but there was no additive effect of anti-PD1. 2’3’-cGAMP alone or co-injected with VLP induced a smaller tumor growth delay and no complete responses were observed. Empty VLP had no effect. Similar trends were observed on mouse survival (defined in this study as the time until the ethical endpoint of 2000 mm^3^ tumor size is reached) (**Figure 1E**). Specifically, anti-PD1 enhanced the survival of mice treated with cGAMP-VLP, while it had no impact when combined with ADU-S100. Furthermore, we observed that the anti-tumor effect of ADU-S100 was characterized by necrosis of all the injected tumors, while necrosis was rarely observed with cGAMP-VLP **(Figure S2A)**.

These results suggested potential differences in T cell responses induced by cGAMP-VLP or ADU-S100. We measured the frequency of OVA-specific CD4^+^ and CD8^+^ T cell responses in blood 10 days after treatment initiation. cGAMP-VLP induced significant responses and the majority of mice showed detectable responses (**Figure 1F**). In contrast, ADU-S100 did not induce detectable T cell responses in most mice. In few mice, a T cell response was detected, but its magnitude did not reach the average response observed with cGAMP-VLP. Overall, the induction of OVA-specific T cell responses by ADU-S100 was not significant. It has been proposed that ADU-S100 ablates the T cell responses, and that at lower doses it may induce tumor-specific T cell responses in blood (Sivick et al., 2018). We performed a dose-titration of ADU-S100 in the B16-OVA model and observed a dose-response anti-tumor effect (**Figure S2B**). In the blood, we detected OVA-specific CD8^+^ responses at the highest dose of ADU-S100 in a subset of mice, but these were not significant (**Figure S2C**). No OVA-specific CD8^+^ response was observed at lower doses of ADU-S100, nor in CD4^+^ T cells. Thus, lower doses of ADU-S100 do not induce tumor-specific T cell responses in blood in this model. We conclude that intra-tumoral injection of cGAMP-VLP stimulates immunogenic anti-tumor T cell responses at low doses of cGAMP.

### Tumor specific T-cell responses elicited by intra-tumorally administered cGAMP-VLP translate into systemic synergy with anti-PD1

We next sought to explore whether the T cell responses induced by cGAMP-VLP translate into systemic anti-tumor effect. To this end, we used a B16-OVA dual tumor model (**Figure 2A**). Intra-tumoral injection of cGAMP-VLP or ADU-S100 in one of the tumors induced IFN-α, IFN-β, IL-6 and TNF-α in the blood (**Figure 2B**). 10 days later, significant levels of OVA-specific CD8+ and CD4+ T cells were detected in the blood of cGAMP-VLP treated mice (**Figure 2C**). In contrast, ADU-S100 induced T cell responses only in a minority of mice that were not statistically significant compared to the control group. We next monitored tumor growth in groups co-treated or not with anti-PD1. We confirmed that B16-OVA was resistant to anti-PD1 (**Figure 2D**). cGAMP-VLP induced a delay in tumor growth in local and distant tumors, and addition of anti-PD1 extended the delay and increased the number of eradicated tumors (**Figure 2D**). In contrast ADU-S100 induced a strong anti-tumor effect that was characterized by necrosis at the injected tumor **(Figure S2D).** At the distal tumor, ADU-S100 induced an anti-tumoral effect, but this effect was not enhanced by anti-PD1 (**Figure 2E**). Ultimately, cGAMP-VLP combined with anti-PD1 decreased the distal tumor size more potently than ADU-S100, irrespectively of its combination with anti-PD1 (**Figure 2F**). Completely responding mice were challenged at day 80 with a second round of tumor graft. Mice that eradicated their initial tumor following cGAMP-VLP treatment were more resistant to the formation of a new tumor than mice that received ADU-S100 (**Figure 2G**). We conclude that cGAMP-VLP demonstrated a synergistic effect with anti-PD1, unlocking the ability of B16-OVA bearing mice to respond to immune checkpoint blockade. In contrast, the synthetic CDN ADU-S100 induces systemic anti-tumor responses that do no elicit OVA-specific T cells response and do not synergize with anti-PD1.

**Figure 2.**
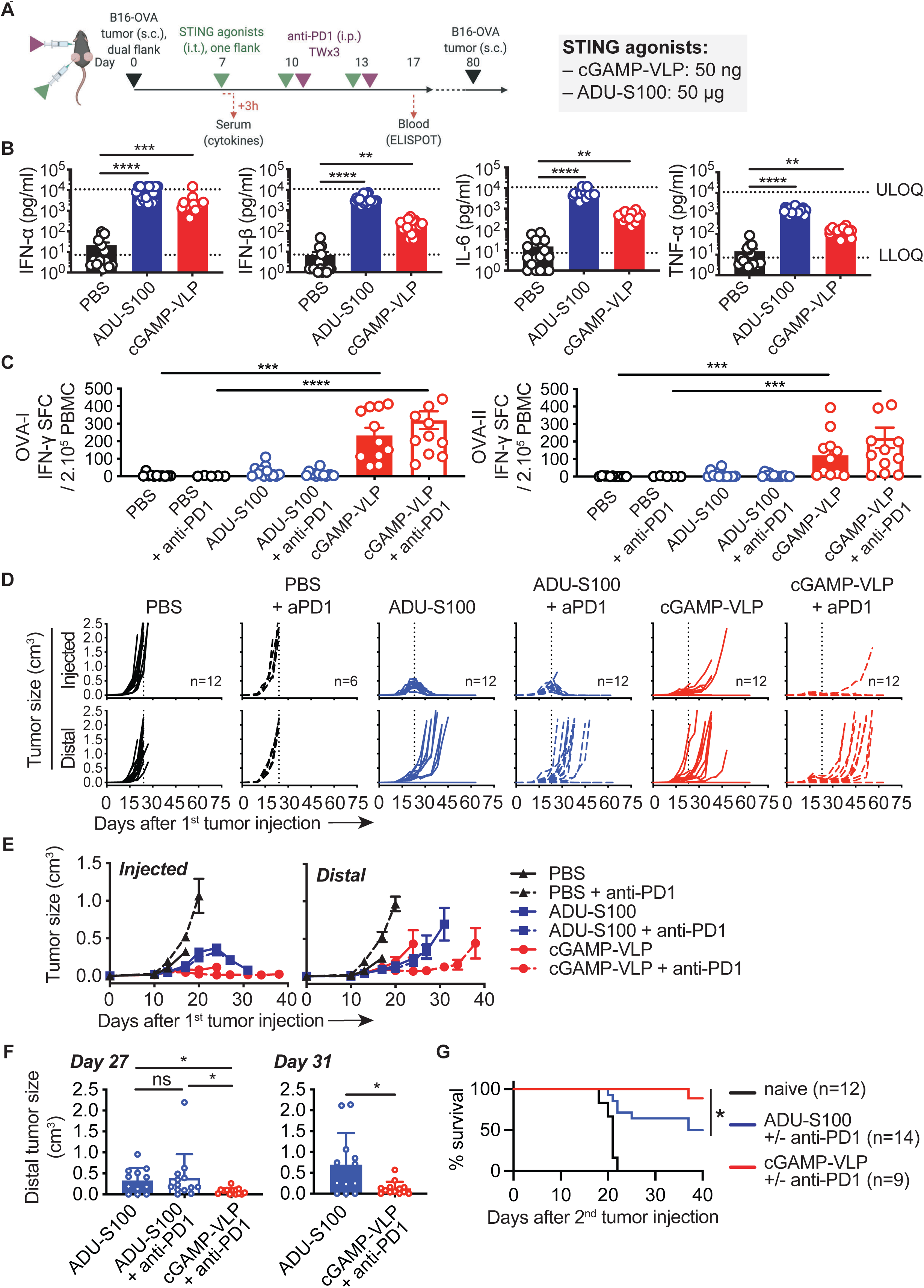
Tumor specific T-cell responses elicited by cGAMP-VLP translate into abscopal synergy with anti-PD1. **(A)** Overview of the experimental design. Treatments were initiated on palpable tumors. **(B)** Concentrations of IFN-α, IFN-β, IL-6 and TNF-α in the serum of B16-OVA dual tumor-bearing mice 3 hours after treatment (bar at mean + SEM, n = 6 to 24 mice per group, combined from 2 independent experiments, Kruskal-Wallis with Dunn post-test, LLOQ = lower limit of quantification, ULOQ = upper limit of quantification). **(C)** Ova-specific CD8 (OVA-I) and CD4 (OVA-II) T cell responses in blood, assess by IFN-γ ELISPOT (bar at mean + SEM, n = 6 to 12 mice per group, combined from 2 independent experiments, Kruskal-Wallis with Dunn post-test). **(D)** Growth curves of individual injected and distal B16-OVA tumors treated as indicated. Vertical dotted line indicates the death of the last mouse in the PBS-injected group. **(E)** Mean growth over time of B16-OVA injected and distal tumors treated as indicated (line at mean + SEM, n = 6 to 12 mice per group, combined from 2 independent experiments). **(F)** Distal tumor size at the indicated days in treated mice, for groups that did not reach ethical limits (line at mean + SEM, n = 12 mice per group, combined from 2 independent experiments, Kurskal-Wallis with Dunn post-test for day 27, Mann-Whitney for day 31). **(G)** Survival of mice after secondary challenge. In complete responding mice, B16-OVA cells were injected 80 days from the first injection of tumor cells and treatments (combined from 3 experiments with single or dual tumors at the first injection, Gehan-Breslow-Wilcoxon test on cGAMP-VLP + anti-PD1 vs ADU-S100 + anti-PD1).

### cGAMP-VLP requires host STING and T cells to induce anti-tumor effects

To understand the nature of the anti-tumor response induced by cGAMP-VLP, we tested the role of STING and T cells using *Sting1* and *Rag2* knock-out mice, respectively (**Figure 3A**). We selected a dual tumor B16-OVA model treated with intra-tumoral 50 ng cGAMP-VLP or 50 µg ADU-S100 monotherapy. Induction of IFN-α, IL-6 and TNF-α by cGAMP-VLP or ADU-S100 was lost in *Sting1^-/-^*mice, indicating that STING is required in host cells (**Figure 3B**). In contrast, the cytokines were still induced in *Rag2^-/-^* mice showing that T cells were not mediating these early response cytokines. Next, we measured the OVA-specific CD4^+^ and CD8^+^ T cell response in blood. As expected, the T cell responses induced by cGAMP-VLP were not detected in *Rag2^-/-^* mice (**Figure 3C**). In *Sting1^-/-^* mice, the T cell responses induced by cGAMP-VLP were heterogeneous and not statistically significant, as compared to WT mice. Nevertheless, T cell responses were detectable in some of the mice, indicating that additional pathways contribute to the immune-stimulating activity of cGAMP-VLP. We next examined the growth of tumors. The anti-tumor effect of cGAMP-VLP and ADU-S100 on the size of injected and distal tumors was lost in *Sting1^-/-^*(**Figure 3D**). In *Rag2^-/-^* mice, the anti-tumor effect cGAMP-VLP was lost in the injected and distal tumors. In contrast, the effect of ADU-S100 was maintained in the injected tumors, but lost at the distal ones. Consistently, cGAMP-VLP and ADU-S100 increased the survival of dual B16-OVA tumor bearing mice compared to PBS treated mice, and these increases were abolished in *Sting1^-/-^* or *Rag2^-/-^* mice (**Figure 3E**).

**Figure 3.**
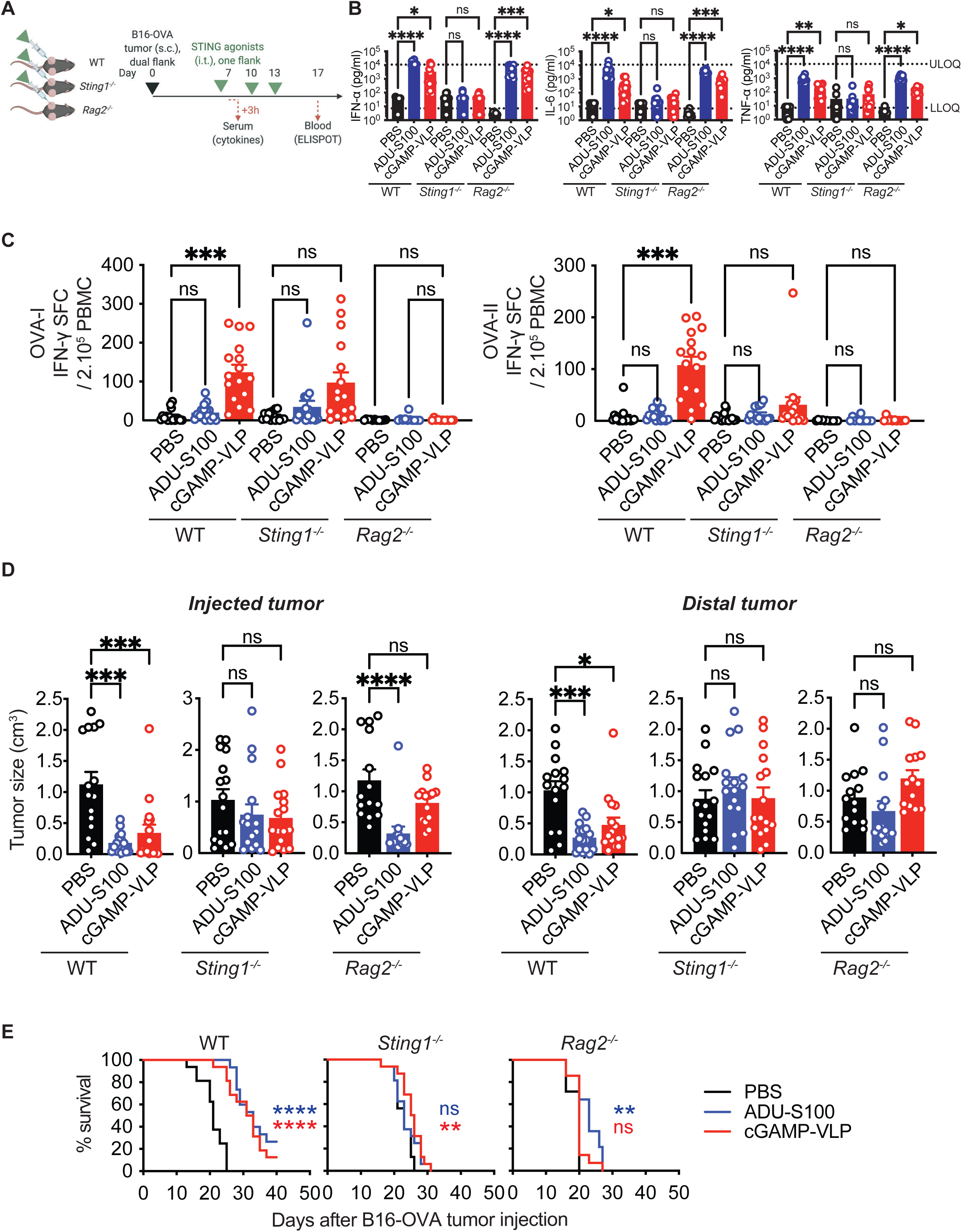
The anti-tumor effect of cGAMP-VLP requires host STING and T lymphocytes. **(A)** Overview of the experimental design using B16-OVA dual tumor-bearing mice (WT, *Sting1^-/-^* or *Rag2^-/-^*). Treatments were initiated on palpable tumors. **(B)** Concentrations of IFN-α, IL-6 and TNF-α in the serum 3 hours after the first treatment by i.t. injection of PBS, 50 µg ADU-S100 or 50 ng cGAMP-VLP (bar at mean + SEM, n = 8 to 16 mice per group, combined from 2 independent experiments, Kruskal-Wallis with Dunn post-test, LLOQ = lower limit of quantification, ULOQ = upper limit of quantification). **(C)** Ova-specific CD8 (OVA-I) and CD4 (OVA-II) T cell responses in blood of WT, *Sting1^-/-^* or *Rag2^-/-^* mice 17 days after tumor implantation, assessed by IFN-γ ELISPOT (bar at mean + SEM, n = 11 to 12 mice per group, combined from 2 independent experiments, Kruskal-Wallis with Dunn post-test). Mice were randomized at day 7 and treated by i.t. injection at days 7, 10 and 13. **(D)** Size of injected and distal tumors 16 days after tumor implantation in WT, *Sting1^-/-^* or *Rag2^-/-^* treated mice (line at mean + SEM, n = 16 mice per group except n = 15 for WT PBS group, combined from 2 independent experiments, Kruskal-Wallis with Dunn post-test). **(E)** Survival of B16-OVA dual tumor-bearing mice (WT, *Sting1^-/-^*or *Rag2^-/-^*) treated as indicated (log-rank Mantel-Cox test).

These results prompted us to test the relative role of CD8^+^ T cells and NK cells in tumor elimination induced by cGAMP-VLP using depleting antibodies (**Figure S3A**). The anti-CD8α antibody induced a depletion of CD8^+^ T cells at day 7 and 17, an increase in NK cells at day 17, and no effect on CD4^+^ T cells (**Figure S3B**). In contrast the anti-NK1.1 antibody depleted NK cells and had a slight depleting effect on CD8^+^ T cells at days 7. The antibodies had no effect on cytokine production induced by cGAMP-VLP at day 7, two days after the first round of depletion (**Figure S3C**). As expected, the anti-CD8α antibody blunted the detection of OVA-specific CD8^+^ T cells (**Figure S3D**). CD8^+^ T cell depletion also cancelled the effect of cGAMP-VLP on mouse survival, while NK cell depletion had no effect (**Figure S3E**). We conclude that the anti-tumor effect of cGAMP-VLP requires STING in the host and CD8^+^ T cells, but not NK cells, while the effect of ADU-S100 requires host STING but is partially independent of T cells.

### Immune cell composition and activation differentiates cGAMP-VLP from ADU-S100

Our results suggested the following paradox: while high levels of tumor-antigen-specific T cells were detected in the blood of cGAMP-VLP treated mice but not in ADU-S100 treated mice, the abscopal anti-tumoral effect of both treatments required T cells. To resolve this paradox, we investigated the composition and activation status of immune cells in tumors and lymphoid organs (**Figure 4A**). In the injected tumors, cGAMP-VLP induced a significant increase in CD8^+^ T cells and a decrease in CD4^+^ Tregs and NK cells (**Figure 4B, top panel**). In contrast, ADU-S100 significantly depleted CD45.2^+^ immune cells, in particular NK and CD4^+^ T cells. ADU-S100 had no impact on CD8^+^ T cells or Tregs. In the distal tumor, cGAMP-VLP induced a significant increase in CD8^+^ T cells but Tregs levels were not affected (**Figure 4B, bottom panel**). In contrast ADU-S100 had no significant impact on the proportion of immune cells based on the markers tested in the distal tumor. We next analyzed lymphoid organs. In the tumor-draining lymph nodes, cGAMP-VLP increased the proportion of effector memory CD4^+^ and CD8^+^ T cells (**Figure 4C, left panel**). In contrast, ADU-S100 decreased the frequency of central memory CD4^+^ T cells, had no impact on effector memory CD4^+^ T cells, and increased the proportion of effector memory CD8^+^ T cells. In non-draining lymph nodes and in the spleen, both cGAMP-VLP and ADU-S100 increased the proportion of effect memory CD8^+^ T cells (**Figure 4C, middle and right panels**). It was surprising that both cGAMP-VLP and ADU-S100 increased effector memory CD8^+^ T cells in all lymphoid organs examined, but only cGAMP-VLP induced robust levels of tumor antigen-specific T cell responses. This raised the possibility that ADU-S100 might induce T cell activation independently from tumor antigens. To test this possibility, we examined the level of CD69, an early marker of T cell activation. Strikingly, ADU-S100 induces upregulation of CD69 in tumors and in all lymphoid organs tested, in both CD4^+^ and CD8^+^ (**Figure 4D**). This reached up to 20% and 30% of T cells in spleen and non-draining lymph nodes, a week after the last injection of ADU-S100. This systemic effect was not observed with cGAMP-VLP, which induced significant levels of CD69 in non-draining lymph nodes, but not in other organs tested. This result suggests that ADU-S100 induces a general activation of T cells, which does not appear to translate into the expansion of tumor antigen-specific T cells. In contrast, cGAMP-VLP appears to induce a specific T cell response for tumor antigens.

**Figure 4.**
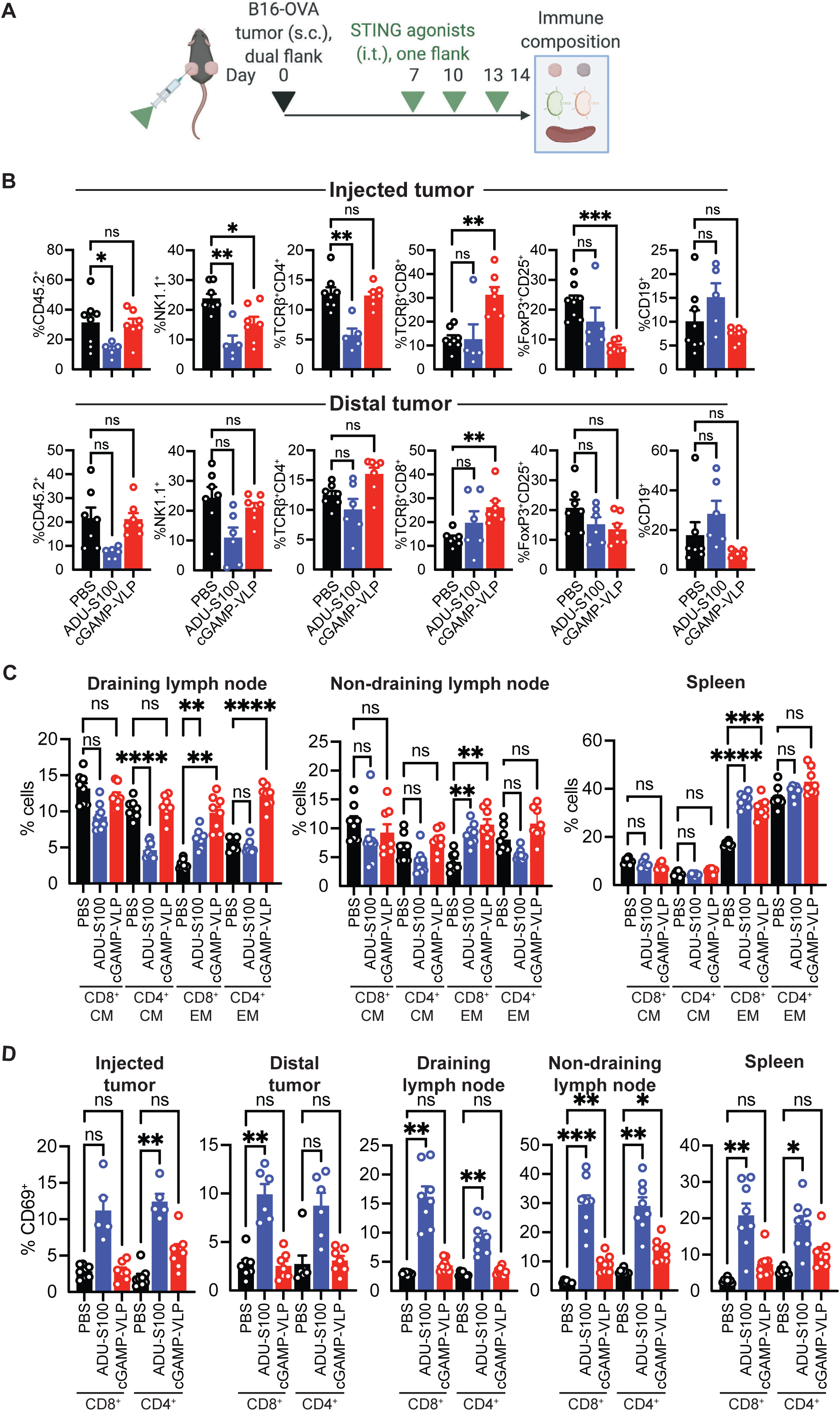
Differential T cell subset composition in response to cGAMP-VLP over ADU-S100. **(A)** Outline of the experiment. **(B)** Frequency of immune cells (%CD45.2^+^ within total live cells), NK cells (%NK1.1^+^ within CD45.2^+^), TCRβ^+^CD4^+^ T cells (within CD45.2^+^), TCRβ^+^CD8^+^ T cells (within CD45.2^+^), Tregs (%FoxP3^+^CD25^+^ within CD45.2^+^TCRβ^+^CD4^+^) and B cells (%CD19^+^ within CD45.2^+^) in B16-OVA dual tumor-bearing mice treated as indicated at days 7, 10 and 13 and analyzed at day 14. Treatments were started on tumors of 10-20 mm3 average volume per group. Data combined from groups with and without anti-PD1 (n=6 to 8 mice per group, Brown-Forsythe and Welch ANOVA test). **(C)** Frequency of central memory (CM, gated as CD44^+^CD62L^+^ within CD45.2^+^TCRβ^+^CD8^+^ or CD4^+^) and effector memory (EM, gated as CD44^+^CD62L^-^ within CD45.2^+^TCRβ^+^CD8^+^ or CD4^+^) T cells in the indicated organs (n=8 mice per group, Brown-Forsythe and Welch ANOVA test). **(D)** Frequency of CD69^+^ cells within CD45.2^+^TCRβ^+^CD8^+^ and CD45.2^+^TCRβ^+^CD4^+^ T cells in the indicated organs (n=6 to 8 mice per group, Brown-Forsythe and Welch ANOVA test).

### cGAMP-VLP targets preferentially antigen-presenting cells

To understand the induction of tumor antigen-specific T cells by cGAMP-VLP, we analyzed its effect *in vitro* on a set of cell types present in the tumor micro-environment, starting with cell lines. We treated the tumor cell line B16-OVA, the endothelial cell line MS1, the dendritic cell line MutuDC and the macrophage cell line RAW. cGAMP-VLP induced the highest levels of IFN-β in RAW cells, followed by MutuDC and MS1, in a dose-dependent manner (**Figure S4A, S4B**). The IFN-β induction in B16-OVA cells was the lowest. ADU-S100 also induced dose-dependent IFN-β, but this was less cell-type selective than cGAMP-VLP. Soluble cGAMP induced detectable IFN-β only at the highest tested dose. To gain further insights in the induction of interferons by antigen-presenting cells, we treated bone marrow derived macrophage (BMDM) and dendritic cells (BMDC), the latter obtained either with GM-CSF (which generates mainly inflammatory dendritic cells) or with FLT3L (which generates a mixed population of cDC1, cDC2 and pDCs). cGAMP-VLP and ADU-S100 induced similar levels of IFN-α and IFN-β in BMDM and BMDC (with GM-CSF) (**Figure S4C**). In contrast, cGAMP-VLP induced significantly higher levels of both cytokines in BMDC (with FLT3L) (**Figure S4D**). Synthetic cGAMP induced detectable cytokines only at the highest tested dose, despite 1000-fold higher amounts than in cGAMP-VLP. These results suggested a preferential activation of STING in antigen-presenting cells by cGAMP-VLP, in particular in FLT3L-derived cells. To determine if this was associated with preferential uptake of the particles, we attempted to detect cGAMP-VLP *in vivo* in samples stained for p24, but the antibody-based detection was not sensitive enough. As a surrogate, we treated splenocytes with cGAMP-VLP and stained for p24 (**Figure S5A**). The highest levels of uptake were detected in macrophages, cDC1 and cDC2 (**Figure S5B, S5C, S5D**). The particles were also detected in some lymphocytes, but only in a fraction of cells within each population. Altogether these results indicate that cGAMP-VLP targets preferentially antigen-presenting cells.

### STING is required in dendritic cells for T-cell mediated anti-tumor effects of cGAMP-VLP

To decipher the contribution of STING within antigen-presenting cells, we generated STING-OST^fl^ mice in which the first coding exon of *Sting1* was flanked by LoxP sites. We also introduced a Twin-Strep-tag (OST) at the N-terminus of STING protein. We crossed the mice to *LysM-cre* or *Itgax-cre* and confirmed preferential deletion of STING in macrophages or dendritic cells, respectively, using Strep-Tactin staining, and thus referred to these mice as STING-OST^ΔMP^ and STING-OST^ΔDC^, respectively (**Figure S6A, S6B**). Following STING deletion in macrophages, the induction of IFN-α and IL-6 in serum by cGAMP-VLP and ADU-S100 was reduced (**Figure 5A, 5B**). However, the induction of OVA-specific T cells by cGAMP-VLP (**Figure 5C**) and the anti-tumoral effect (**Figure 5D**) were maintained. In comparison, the anti-tumor effect of ADU-S100 was partially reduced. Following STING deletion in dendritic cells, the induction of IFN-α and IL-6 by cGAMP-VLP was reduced, but not for ADU-S100 (**Figure 5E**). The induction of OVA-specific T cells by cGAMP-VLP was reduced, but not completely lost (**Figure 5F**) and the anti-tumor effect of cGAMP-VLP was essentially abrogated in these mice (**Figure 5G**). In contrast, the anti-tumor effect of ADU-S100 was reduced but maintained. These results indicate that STING is specifically required in dendritic cells for the anti-tumor effect of cGAMP-VLP, while the anti-tumor effect of ADU-S100 depends partially on STING in macrophages and dendritic cells.

**Figure 5.**
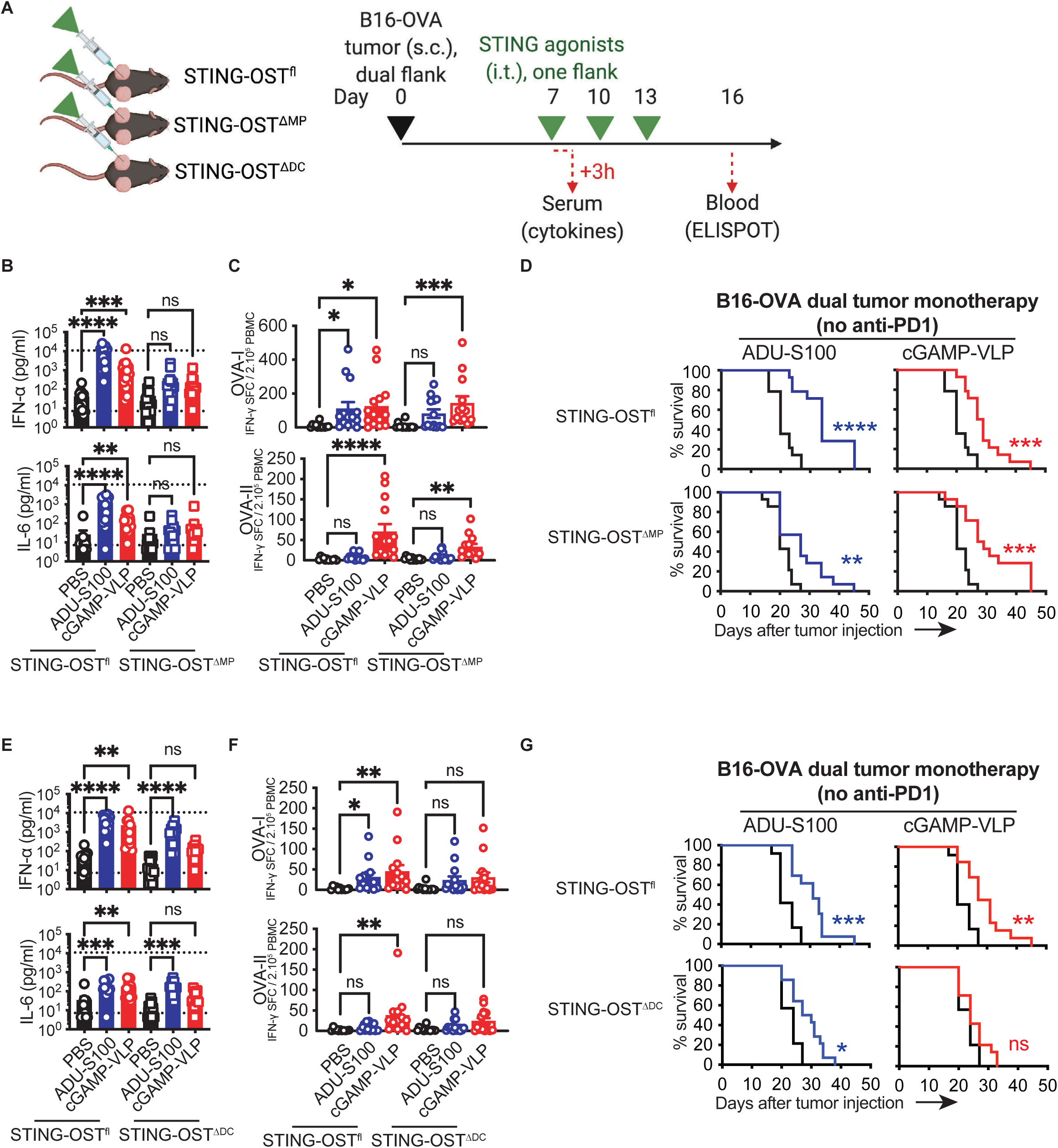
Anti-tumor effect of cGAMP-VLP requires STING in dendritic cells. **(A)** Outline of the experiment using B16-OVA dual tumor-bearing mice (STING-OST^fl^, STING-OST^ΔMP^ or STING-OST^ΔDC^). Treatments were initiated on palpable tumors **(B)** Concentrations of IFN-α and IL-6 in the serum of STING-OST^fl^ or STING-OST^ΔMP^ mice 3 hours after the first treatment by i.t. injection of PBS, 50 µg ADU-S100 or 50 ng cGAMP-VLP (bar at mean + SEM, n = 14 mice per group, combined from 2 independent experiments, Kruskal-Wallis with Dunn post-test, LLOQ = lower limit of quantification, ULOQ = upper limit of quantification). **(C)** Ova-specific CD8 (OVA-I) and CD4 (OVA-II) T cell responses in blood of STING-OST^fl^ or STING-OST^ΔMP^ mice treated as indicated, 16 days after tumor implantation, assessed by IFN-γ ELISPOT (bar at mean + SEM, n = 12 to 14 mice per group combined from 2 independent experiments). **(D)** Survival of B16-OVA dual tumor-bearing STING-OST^fl^ or STING-OST^ΔMP^ mice treated as indicated (n = 14 mice per group combined from 2 independent experiments, log-rank Mantel-Cox test). **(E)** Concentrations of IFN-α and IL-6 in the serum of STING-OST^fl^ or STING-OST^ΔDC^ mice 3 hours after the first treatment by i.t. injection of PBS, 50 µg ADU-S100 or 50 ng cGAMP-VLP (bar at mean + SEM, n = 12 to 14 mice per group, combined from 2 independent experiments, Kruskal-Wallis with Dunn post-test, LLOQ = lower limit of quantification, ULOQ = upper limit of quantification). **(F)** Ova-specific CD8 (OVA-I) and CD4 (OVA-II) T cell responses in blood of STING-OST^fl^ or STING-OST^ΔDC^ mice treated as indicated, 16 days after tumor implantation, assessed by IFN-γ ELISPOT (bar at mean + SEM, n = 11 to 14 mice per group combined from 2 independent experiments). **(G)** Survival of B16-OVA dual tumor-bearing STING-OST^fl^ or STING-OST^ΔDC^ mice treated as indicated (n = 12 to 14 mice per group combined from 2 independent experiments, log-rank Mantel-Cox test).

### Systemic administration of cGAMP-VLP activates anti-tumor T cells immunity

The activation of STING in dendritic cells by cGAMP-VLP raised the possibility that it could induce anti-tumor T cell responses even after injection outside of the tumor mass. We first tested the B16-OVA model combined with anti-PD1 (**Figure 6A**). Sub-cutaneous (s.c.) injection of cGAMP-VLP induced detectable levels of IFN-α, IFN-β, IL-6 and TNF-α, albeit to lower levels than following intra-tumoral (i.t.) injection (**Figure 6B**). Tumor growth was delayed after s.c. injection of cGAMP-VLP (**Figure 6C**), leading to significantly smaller tumors (**Figure 6D**). cGAMP-VLP s.c. also induced anti-OVA T cell responses (**Figure 6E**) and increased the survival of tumor-bearing mice (**Figure 6F**).

**Figure 6.**
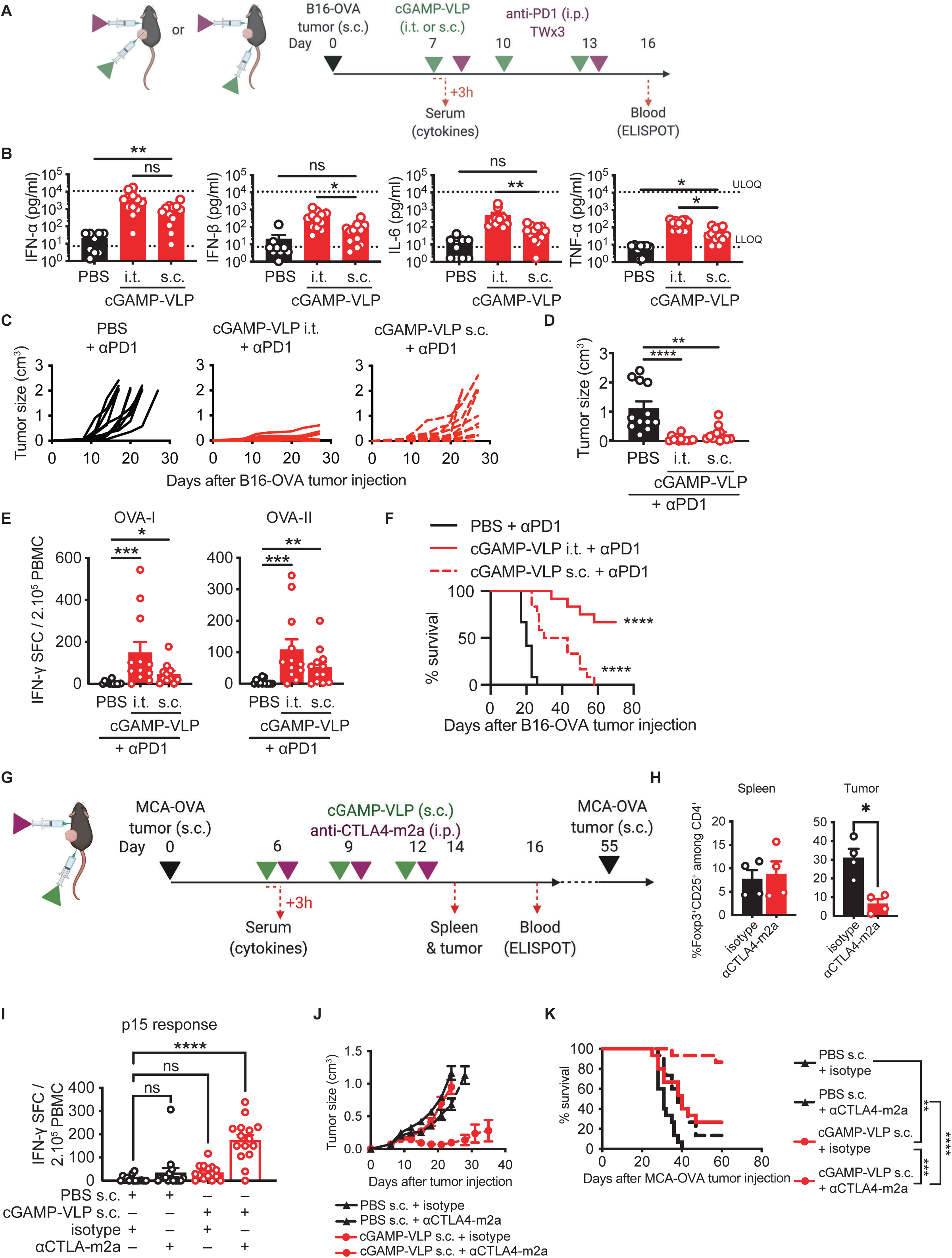
Sub-cutaneous injection of cGAMP-VLP induces anti-tumor synergy with tumor Treg depletion. **(A)** Outline of the experiment using B16-OVA tumors to compare i.t. and s.c. injection routes of cGAMP-VLP. Treatments were started on tumors of 50 mm^3^ average volume per group. **(B)** Concentrations of IFN-α, IFN-β, IL-6 and TNF-α in the serum of mice 3 hours after the first treatment with PBS or 50 ng cGAMP-VLP injected by the i.t. or s.c. route (bar at mean + SEM, n = 9 to 11 mice per group, combined from 2 independent experiments, Kruskal-Wallis with Dunn post-test, LLOQ = lower limit of quantification, ULOQ = upper limit of quantification). **(C)** Growth curves of individual B16-OVA tumors in mice treated as indicated (n = 18 mice per group combined from 3 independent experiments). Mice were randomized at day 7, and treated at days 7, 10 and 13 with cGAMP-VLP, and bi-weekly from day 7 for 3 weeks with anti-PD1. **(D)** Size of tumor 17 days after tumor implantation in treated mice (line at mean + SEM, n = 18 mice per group combined from 3 independent experiments, Kruskal-Wallis with Dunn post-test). **(E)** Ova-specific CD8 (OVA-I) and CD4 (OVA-II) T cell responses in blood of mice 16 days after tumor implantation, assessed by IFN-γ ELISPOT (bar at mean + SEM, n = 12 mice per group, combined from 2 independent experiments, Kruskal-Wallis with Dunn post-test). **(F)** Survival of B16-OVA tumor-bearing mice treated as indicated (log-rank Mantel-Cox test, n = 12 mice per group combined from 2 independent experiments). **(G)** Outline of the experiment using MCA-OVA tumors, cGAMP-VLP and a tumor Treg-depleting antibody (anti-CTLA4-m2a). Treatments were started on tumors of 50 mm^3^ average volume per group. **(H)** Fraction of CD25^+^FoxP3^+^ Tregs within CD45.2^+^TCRβ^+^CD4^+^ cells in spleen and tumor, 48 hours after last i.p. injection of αCTLA4-m2a or isotype (n=4, 2 mice from 2 independent experiments were analyzed). **(I)** CD8 T cell responses against p15 antigen in blood of mice 16 days after tumor implantation, assessed by IFN-γ ELISPOT (bar at mean + SEM, n = 15 mice per group, combined from 2 independent experiments, Kruskal-Wallis with Dunn post-test). **(J)** Mean growth over time of MCA-OVA tumors treated as indicated (line at mean + SEM, n = 15 mice per group, combined from 2 independent experiments). **(K)** Survival of MCA-OVA tumor-bearing mice treated as indicated (n = 15 mice per group, combined from 2 independent experiments, log-rank Mantel-Cox test).

In these experiments, the i.t. route remained more effective than the s.c. route at inducing T cell responses and anti-tumor effects. This suggested that a negative regulator of the immune response might be eliminated locally by i.t. activation of STING. We previously noted that cGAMP-VLP induced a reduction of Tregs in the injected tumor, but not in the distal tumors (**Figure 4B**). This raised the possibility that intra-tumor Tregs might limit the anti-tumor effect of systemic STING activation by cGAMP. In order to test this hypothesis, we used an IgG2a isotype antibody against CTLA4 (anti-CTLA4-m2a), which has been shown to selectively deplete Tregs in tumors (Arce Vargas et al., 2017; Selby et al., 2013), and we confirmed this effect in the MCA-OVA tumor model (**Figure 6G, 6H**). Treatment with anti-CTLA4-m2a had no effect on the induction IFN-α, IL-6 and TNF-α by cGAMP-VLP (**Figure S7A**). In monotherapy, cGAMP-VLP s.c. or anti-CTLA4-m2a increased the frequency of OVA-specific CD8^+^ and CD4^+^ T cells, but no significant response to the endogenous tumor antigen p15 (**Figure 6I, S7B**). In contrast, combining cGAMP-VLP s.c. with anti-CTLA4-m2a synergized to significantly increase the levels of T cells against p15, and further increased the levels of T cells against OVA. Accordingly, combination therapy induced a near-complete reduction in tumor size (**Figure 6J, S7C**). Similarly, monotherapies induced an increase in survival, but only the combination therapy induced long-term survival of treated mice (**Figure 6K**). Completely responding mice were also protected from a secondary tumor challenge (**Figure S7D**). We conclude that systemic administration of cGAMP-VLP activates anti-tumor T cell immunity that synergizes with tumor Treg depletion.

## Discussion

These results highlight the crucial importance of targeting STING activation in particular cell types, namely dendritic cells, to optimize the antigen-specific anti-tumor responses. STING was previously shown to be required in dendritic cells *in vitro* to induce an interferon response to immunogenic tumor cells or tumor DNA (Deng et al., 2014; Woo et al., 2014). *In vivo*, it was previously noted that dendritic cells are a major source of IFN-β in tumors that induce STING-dependent immunogenic responses (Andzinski et al., 2016). Intriguingly, STING in CD11c^+^ cells is also implicated in the negative regulation of allogeneic responses (Wu et al., 2021). Altogether, STING in dendritic cells emerges as a linchpin for the induction of antigen-specific T cell responses.

In contrast to cGAMP-VLP, the anti-tumor responses induced by ADU-S100 were not associated with the induction of tumor-specific T cells. It was previously proposed that the induction of antigen-specific T cells by ADU-S100 was dose-dependent (Sivick et al., 2018). We did not observe such bimodal behavior in the tumor model we tested. We noted that ADU-S100 induced some level of tumor-specific T cells in experiments with in-house bred mice (**Figures 5C, 5F**), but not with mice obtained from an external source (**Figures 1F, 2C**, **S2C**). This raises the intriguing possibility that housing parameters such as the composition of the microbiota, or genetic background, might affect the immunogenic properties of synthetic CDNs. We also noted that synthetic CDNs induced necrosis at the intra-tumoral injection site which was rarely seen with cGAMP-VLP. This is consistent with a role of STING activation in endothelial cells caused by synthetic CDNs as contributing to its local anti-tumor effects (Demaria et al., 2015; Francica et al., 2018; Jeong et al., 2021). The reduced dose of cGAMP in cGAMP-VLP compared to free CDN likely contributes to the reduced tissue necrosis after cGAMP-VLP treatment.

Multiple approaches have been proposed to optimize delivery of CDNs for use as immunomodulators in the absence of exogenous tumor antigens. Synthetic nanoparticles assembled in the presence of CDNs have been shown to enhance cytosolic delivery and activate STING-dependent anti-tumor responses (Lu et al., 2020; Wilson et al., 2018). Exosomes loaded with CDNs appear to achieve similar enhancements (Jang et al., 2021; McAndrews et al., 2021). Principles to ensure that delivery with synthetic approaches will yield tumor-specific T cell responses generated are ill-defined. A common limitation of synthetic cargos and exosomes lies in the passive delivery mechanism to target cells. In contrast, cGAMP-VLPs employ a viral fusion glycoprotein to efficiently fuse with target cells. The size of the VLPs, their lipid bilayer originating from a producer cell and the fusion triggered by VSV-G in acidic endosomes most likely contribute to the selectivity of cGAMP-VLPs for antigen-presenting cells, in particular dendritic cells. Accordingly, retroviral particles are also efficiently captured by antigen-presenting cells *in vivo* (Sewald et al., 2015). In addition, a higher expression of STING or downstream signaling proteins in antigen-presenting cells might also contribute.

A feature of the response to cGAMP-VLP is the decrease of tumor Tregs when it was directly injected in the tumor. We do not know if this effect is a response of Tregs to STING activation in the tumor micro-environment, or whether it is a secondary effect resulting from anti-tumor T cell stimulation. Similar to previous studies, we found that treatment with anti-CTLA4-m2a induced a partial anti-tumor response (Arce Vargas et al., 2017). Combination of s.c. cGAMP-VLP with this tumor Treg-depleting agent induced a near-complete response to treatment. These results suggest that the level of Tregs in the tumor may be an important factor to consider for clinical development of STING-targeted therapies such as cGAMP-VLP.

Altogether, our results establish that cell-type specific activation of STING plays a critical role in anti-tumor immunogenicity. Synthetic STING agonists appear to induce promiscuous STING activation that does not necessarily entail priming of tumor-specific T cells. In contrast, cGAMP-VLP constitutes a biological product that activates STING preferentially in dendritic cells, leading to activation of tumor-specific T cells, which synergize with ICB and Treg depletion. Biological stimulation of STING with cGAMP-VLP has the potential, similar to other biological drugs such as antibodies and CAR-T cells, to contribute to a meaningful treatment regimen to induce anti-tumor immune responses in patients.

## Acknowledgements

We thank S. Amigorena, J. Rehwinkel and N. de Silva for discussions, X. Lahaye and C. Conrad for help with experiments. This work supported by Stimunity, Institut Curie, Fondation Carnot, INSERM, Association Nationale de la Recherche et de la Technologie (Cifre to A.B.), European Research Council (ERC-2016-PoC STIMUNITY), Fondation BMS, Cancéropôle Ile-de-France (STINGTARGET), Agence Nationale de La Recherche (LABEX DCBIOL, ANR-10-IDEX-0001-02 PSL, ANR-11-LABX-0043), the MSDAVENIR Fund (to B.M.), and the Investissement d’Avenir program PHENOMIN (French National Infrastructure for mouse Phenogenomics; ANR10-INBS-07 to BM). Request for STING-OST^fl^ mice should be addressed to B. Malissen.

## Author contributions

B. Jneid performed most experiments, analyzed data and prepared figures. A. Bochnakian performed a set of in vitro experiments. F. Delisle, E. Djacoto and J. Denizeau contributed to experiments. C. Sedlik, R. Kramer, I. Walters, E. Piaggio suggested experiments and contributed to data analysis and interpretations. B. Malissen and F. Fiore conceived and developed the STING-OST^fl^ mice. S. Carlioz developed Randmice. S. Carlioz and N. Manel conceived the study. N. Manel and B. Jneid wrote the manuscript.

## Methods

### Cell culture

293T cells, RAW cells and MS1 cells were cultured in DMEM GlutMAX, 10% fetal bovine serum (FBS) (Gibco), and penicillin-streptomycin (Gibco). THP-1 cells were cultured in RPMI GlutMAX medium, 10% FBS (Gibco), and penicillin-streptomycin (Gibco). B16-OVA cells were cultured in RPMI GlutMAX medium with 10% FBS (Gibco), penicillin-streptomycin (Gibco), 1 mM 2-mercaptoethanol, geneticin and hygromycin. MCA-OVA cells were cultured in RPMI GlutMAX medium with 10% FBS (Gibco), penicillin-streptomycin (Gibco), 1 mM 2-mercaptoethanol, and hygromycin. MB49 cells were cultured in DMEM GlutMAX medium with 10% FBS (Gibco) and penicillin-streptomycin (Gibco). MutuDC were cultured as described (Kozik et al., 2020). The splenocytes were culture in RPMI GlutMAX with 10% FBS (Gibco), penicillin-streptomycin (Gibco), 1 mM 2-mercaptoethanol.

### Cell differentiation from bone marrow

Femurs, shin and fibula of female mice were collected immediately after sacrifice, the fat and muscle tissues were removed, the end of the bones were cut with a pair of scissors, and put in a 0.5 mL tubes in which holes were made at the bottom with a needle. The 0.5 mL tube was put in a 1.5 mL tube containing 200 μL of complete IMDM (Iscove’s modified Dulbecco’s medium, 10% FBS, penicillin-streptomycin a 1mM 2-mercaptoethanol), and centrifugated at 11,000g for 10 seconds.

For BMDM cells were seeded at the concentration of 1 million cells per mL in 20 mL total, in a 20 cm non-tissue culture treated plates in BMDM culture media (RPMI GlutMAX, 10% FBS, penicillin-streptomycin, 1 mM 2-mercaptoethanol, 1 mM sodium pyruvate, non-essential amino acids, HEPES, 10ng/mL human M-CSF (Miltenyi Biotec). Adherent cells were detached with 5 mM EDTA in PBS at day 6. Differentiation was analyzed by staining with anti-CD11b and anti-F4/80 followed by cytometry analysis.

For BMDC (GMCSF), cells were plated in 20 cm non-tissue culture treated plates, at a concentration of 1 million cells per mL in 20 mL, in IMDM containing conditioned supernatant from J558 cells as described (Alloatti et al., 2016). At day 4, non-adherent cells were collected, and loosely adherent cells were collected with 5 mM EDTA in PBS. Non-adherent and loosely adherent cells were combined and seeded at the concentration of 0.5 million cells per mL in 20 mL. At day 7, non-adherent cells were discarded, loosely adherent cells were collected with PBS-EDTA and replated at concentration of 0.5 million cells per mL in 20 mL. At day 10 non-adherent cells were discarded, loosely adherent cells were collected with PBS-EDTA. Differentiation was analyzed by staining with anti-CD11b and anti-CD11c followed by cytometry.

For BMDC (FLT3L), bone marrow was isolated as described above, and plated in 6-well cell culture plates at the concentration of 1.5 million cells per mL in 4mL total of complete IMDM medium supplemented with FLT3L (200ng/ml, Peprotech). At day 10 of differentiation, the loosely adherent cells were harvested using PBS/EDTA and differentiation was checked by staining for MHC-II, CD11c, B220, and CD24.

### cGAMP-VLP production for *in vivo* use

7.5 million 293T cells were plated in 150cm² cell culture flask and incubated overnight. One batch of cGAMP-VLP was made from 4 flasks. The following day, each flask was transfected with 13 μg of pVAX1-cGAS, 8.1 μg of HIV-1 psPAX2, 3.3 μg of pVAX1-VSVG-INDIANA2, and 50 μL of PEIpro (Ozyme reference POL115-010), according to the manufacturer’s instructions. The transfection mixes were prepared in Opti-MEM (Gibco). The morning following transfection, the medium was changed with 52 mL of warm VLP production medium (293T culture medium with 10 mM HEPES and 50 μg/mL Gentamicin). One day later, the cGAMP-VLP-containing supernatant was harvested from the cells, centrifuged for 10 minutes at 200 g 4°C, and filtered through 0.45 µm nylon mesh filters (Fisher 22363547). 39 mL of cGAMP-VLP-containing supernatant was gently overlaid on 6 mL of cold PBS containing 20% sterile filtered endotoxin free sucrose in 6 Ultra-Clear tubes (Beckman Coulter, ref 344058), and centrifuged for 1 hour and 30 minutes at 100,000 g 4°C. The liquid phase was gently aspirated, the pellets were resuspended in cold PBS and transferred to one Ultra-Clear 13.2 mL tube (Beckman Coulter, ref 344059) and centrifuged again at 100,000g 4°C for 1 hour and 30 minutes. The PBS was gently poured out and the pellet was resuspended in 320 µl of cold PBS. Batches were split in 3 aliquotes of 100 µL for experimental use. The remaining 20 µL were diluted 1:4 with 60 µL of PBS and split in 8 aliquotes of 10 µL for quality control assays. Aliquotes were stored at -80°C.

### cGAMP quantification

2’3’-cGAMP ELISA Kit (Cayman Chemical) was used for the quantification of cGAMP in cGAMP-VLP according to the manufacturer’s instructions. After performing the assay, the plate was read at a wavelength of 450 nm. Data was fitted to a 4-parameter sigmoidal curve.

### Biological activity assay of cGAMP-VLP

50,000 THP-1 cells were plated in round bottom 96 well plates in 100 μLof medium, and stimulated with serial-dilutions of cGAMP-VLPs, soluble cGAMP or soluble ADU-S100 in 100 µl. Where indicated, CDN (6 µg) were mixed with lipofectamine 2000 (6 µl) in Opti-MEM (12.75 µl each) following manufacturer’s instructions. The cells were incubated for 18 to 24 hours and stained with an anti-human SIGLEC-1 (Miltenyi ref 130-098-645), fixed in PFA 1% and acquired using a BD FACSVerse cytometer.

### Electron microscopy

cGAMP-VLP suspension was deposited on formvar/carbon–coated copper/palladium grids before uranyl/acetate contrasting and methyl-cellulose embedding for whole-mount. Images were acquired with a digital camera Quemesa (EMSIS GmbH, Mu nster, Germany) mounted on a Tecnai Spirit transmission electron microscope (FEI Company) operated at 80kV.

### Nanoparticle Tracking Analysis

The cGAMP-VLPs were serially diluted in PBS at room temperature and acquired on a NanoSight as previously described (Liao et al., 2019).

### Mice

All animals were used according to protocols approved by Animal Committee of Curie Institute CEEA-IC #118 and maintained in pathogen-free conditions in a barrier facility. Experimental procedures were approved by the Ministère de l’enseignement supérieur, de la recherche et de l’innovation (APAFIS#11561-2017092811134940-v2) in compliance with the international guidelines. C57BL/6J mice were purchased from Charles River Laboratories. C57BL/6J *Rag2^tm1.1Cgn^* (*Rag2^-/-^)* mice were maintained at Centre d’Exploration et de Recherche Fonctionnelle Expérimentale. C57BL/6J *Lyz2^tm1(cre)Ifo^* (LysM-cre), C57BL/6J Tg(Itgax-cre)1-1Reiz (Cd11c-cre), C57BL/6J *Sting^gt/gt^ (Sting1^-/-^)* and STING-OST^fl^ mice were maintained at Institut Curie Specific Pathogen Free facility. Mice were allowed to acclimate to the experimental housing facility for at least three days before tumor injections.

### Generation of STING-OST^fl^ knock-in mice

The mouse *Sting1* gene (also called Tmem173; ENSMUSG00000024349) was edited using a double-stranded HDR template (targeting vector) containing 867 and 1260 bp-long 5’ and 3’ homology arms, respectively. It included a loxP site and a frt-neo^r^-frt cassette that were both inserted in intron 2, 110 bp upstream of the start codon, a Twin-Strep-tag-coding sequence (OST; (Junttila et al., 2005)) that was appended at the 5’ end of the first coding exon (exon 3), and a loxP site located in intron 3, 40 bp downstream of the 3’ end of exon 3. The final targeting vector was abutted to a cassette coding for the diphtheria toxin fragment A (Soriano, 1997). Two sgRNA-containing pX330 plasmids (pSpCas9; Addgene, plasmid ID 42230) were constructed. In the first plasmid, two sgRNA-specifying oligonucleotide sequences (5’-CACCGAGTAGCCCATGGGACTAGC-3’ and 5’-AAACGCTAGTCCCATGGGCTACTC-3’) were annealed, generating overhangs for ligation into the BbsI site of plasmid pX330. In the second plasmid, two sgRNA-specifying oligonucleotide sequences (5’-CACCGTCAAGGGTGTGATACTTGC-3’ and 5’-AAAC-GCAAGTATCACACCCTTGAC-3’) were annealed and cloned into the BbsI site of plasmid pX330. The protospacer-adjacent motifs (PAM) corresponding to each sgRNA and present in the targeting vector were destroyed via silent mutations to prevent CRISPR-Cas9 cleavage. JM8.F6 C57BL/6N ES cells (Pettitt et al., 2009) were electroporated with 20 µg of targeting vector and 2.5 µg of each sgRNA-containing pX330 plasmid. After selection in G418, ES cell clones were screened for proper homologous recombination by Southern blot and PCR analysis. A neomycin specific probe was used to ensure that adventitious non-homologous recombination events had not occurred in the selected ES clones. Mutant ES cells were injected into BalbC/N blastocysts. Following germline transmission, excision of the frt-neo^r^-frt cassette was achieved through genetic cross with transgenic mice expressing a FLP recombinase under the control of the actin promoter (Rodríguez et al., 2000). Two pairs of primers were used to distinguish the WT and edited *Tmem173* alleles. A pair of primers (sense 5’-TGTAGGATGCTATGTGCCCA-3’ and antisense 5’-GATCCCAGCCCAACTCAGCT-3’) amplified a 501 bp-long band in the case of the wild-type *Tmem173* allele and a 722 bp-long band in the case of the mutant allele.

The resulting STING-OST^fl^ mice (official name B6-*Tmem173*^Tm1Ciphe^ mice) have been established on a C57BL/6N background. They express a multitask *Tmem173* allele in which the third exon of the *Tmem173* allele is bracketed by *loxP* sequences and a sequence corresponding to an affinity Twin-Strep-Tag (OST) is appended at the 5’ end of the ORF of the *Tmem173* gene. When bred to mice that express tissue-specific Cre recombinase, the resulting offspring will have exon 3 removed in the Cre-expressing tissues, resulting in cells lacking STING.

### Mouse randomization

Mouse randomizations were performed using Randmice (https://randmice.com) based on tumor volume to distribute mice and homogenize the average tumor volume within the different groups. The algorithm randomly shuffles all mice between the groups and calculates the average tumor volume for each group. 10e9 iterations are performed in order to minimize the difference in tumor volume average between all groups.

### Tumor implantation

Female mice were inoculated subcutaneously on the lower right or right and left flanks with 5×10^5^ B16-OVA cells in 100 µL of HBSS or with 5×10^5^ MB49 cells in 100 µL of PBS . Mice were monitored for morbidity and mortality daily. Tumors were monitored twice or three times per week. Mice were euthanized if ulceration occurred or when tumor volume reached 2000 mm^3^. Tumor sizes were measured using a digital caliper and tumor volumes calculated with the formula (length x width^2^)/2. Following tumor implantation, mice were randomized into treatment groups using the Randmice software. In some experiments, tumor-free survivors were challenged with tumor cells on the opposite, non-injected flank several weeks after the collapse of the primary tumor. Naive mice of the same age were used as controls.

### *In vivo* immunotherapy

Intra-tumoral (i.t.) or subcutaneous (s.c.) injections were initiated when tumors are palpable or reached close to 50 mm3 (40-80 mm^3^), as indicated in legends. A U-100 insulin syringe or equivalent [0.33 mm (29 G) x 12.7 mm (0.5 mL)] was filled with 50 µl of samples (VLP, cGAMP-VLP or synthetic CDN diluted in PBS) and all air bubbles were removed. Mice were anesthetized with isoflurane. With the bevel facing the skin, the needle was injected shallowly into the area directly adjacent to the tumor, and the needle was moved underneath the skin until it reached the inside back of the tumor. The samples were injected slowly into the center of the tumor (for the i.t.) or under the skin, 1 cm from the border of the tumor (for the s.c.). The needle was then removed delicately to avoid reflux. Treatments consisting of 200 µg of αPD1 antibody (clone RMP1-14, BioXcell) or 200 µg isotype control antibody (Rat IgG2a, BioXcell) were diluted in PBS at 1 mg/ml and administered by intra-peritoneal (i.p.) injection at the indicated time points.

### *In vivo* antibody depletion

For CD8^+^ and NK1.1 depletions studies, B16-OVA tumor bearing mice were treated with 200 µg of anti-CD8α monoclonal antibody (clone 53-6.7, BioXcell) or 200 µg of anti-NK1.1 monoclonal antibody (clone PK136, BioXcell) or 200 µg of isotype control antibody (Rat IgG2a, BioXcell) two times prior and four times after i.t. treatment with STING agonists. To confirm the cell depletion, PBMC were stained according to standard protocols before depletion, at day 7 and day 17. Briefly, cells were surface-stained in 100 µL antibody-mix in FACS buffer: CD19 (clone 6D5), TCR-b (clone H57-597), CD4 (clone RM4-5), CD8 (Life Technologies) and NK1.1 (clone PK136). For Treg (Foxp3+CD25+ cells) depletion, MCA-OVA tumor bearing mice were treated with 200 µg of anti-mCTLA4-mIgG2a monoclonal antibody (Invivogen) or 200 µg of isotype control antibody (mouse IgG2a, Invivogen) three times at days 6, 9 and 12 after tumor engraftment. To confirm the Treg depletion, spleen and tumor cells were stained according to standard protocols 48 hours after the last antibody injection. Briefly, cells were surface-stained in 100 µL antibody-mix in FACS buffer: CD45.2 (clone 104), CD19 (clone 6D5), TCR-b (clone H57-597), CD4 (clone RM4-5), CD8 (Life Technologies) and CD25 (clone PC61), followed by an intracellular staining in 50 µL with anti-Foxp3 (clone FJK-16s) and anti-Ki67 (BD Biosciences).

### ELISPOT Assay

T cell responses were assessed by IFN-γ ELISPOT 10 days after the first i.t. injection of cGAMP-VLP, synthetic CDNs or PBS. Mice were bled from the retro-orbital sinus. PBMCs were isolated from whole blood by lysing the red blood cells with an ammonium chloride lysis buffer (NH_4_Cl 1.5 M, NaHCO_3_ 100 mM, EDTA 10 mM). 2×10^5^ PBMCs were plated per well in the RPMI medium containing 10% FBS and 1% penicillin-streptomycin. PBMCs were stimulated overnight with media as a negative control, Dynabeads mouse T-activator CD3/CD28 (GIBCO) as a positive control, 10 µg/mL OVA-I 257-264 peptide (SIINFEKL) or 40 µg/mL OVA-II 265-280 peptide (TEWTSSNVMEERKIKV) or 10 µg/mL p15E peptide (KSPWFTTL) or 10 µg/mL DBy 608-622 peptide (NAGFNSNRANSSRSS) or 10 µg/mL UTy 246-254 (WMHHNMDLI). Spots were developed using mouse IFN-γ ELISPOT antibody pair (Diaclone) according to the manufacturer’s instructions. The number of spots was enumerated using an ImmunoSpot analyzer and evaluated by subtracting the specific values from the negative control spot number of each sample.

### Stimulation of cells with CDNs and cGAMP-VLP

100,000 of the indicated cells were seeded in flat bottom 96-well plates in 200 μL and incubated for few hours until attached to the plate. 100 μL were removed and replaced with serial dilutions of ADU-S100, cGAMP, cGAMP-VLP or empty VLP. Cells were incubated for 18 hours, and IFN-α and IFN-β were measured in the supernatant.

### cGAMP-VLP capture by splenocytes *in vitro*

Spleens were harvested from female C57BL6/J mice. Splenocytes were isolated by pressing the organ through a 40 µm cell strainer. Red blood cells were lysed using an ammonium chloride lysis buffer as described above. 1 to 3 million cells were plated in a 96-well round bottom plate in 150 μL of medium. 50μL of cGAMP-VLP or PBS was added and cells were incubated overnight at 37°C 5% CO_2_. The following day the cells were stained with antibodies against extracellular markers (MHC-II eFluor450, eBioscience 48-5321-82; CD4 BV785, bioLegend 100552; NK1.1 PerCP-Cy5.5, BD Biosciences 561111; CD11b PE, Invitrogen 12-0112-82; CD11c PETR, Invitrogen MCD11c17; CD19 PE-Cy5, Invitrogen 15-0193-82; TCR-β PE-Cy7, bioLegend 109222; CD8 APC, BD biosciences 561093; F4/80 AF700, eBioscience 56-4801-82; Fixable Viability Dye, eFluor780; eBioscience 65-0865-14), washed and permeabilized using the BD Cytofix/Cytoperm Fixation Permeabilization Solution kit (reference 554714) according to the manufacturer’s instructions. The cells were then washed with the permeabilization buffer, following by staining for 15 minutes at room temperature with a 1:100 dilution of a fluorescent anti-HIV-1 GAG antibody (KC57-FITC, Beckman Coulter reference 6604665) in permeabilization buffer. Cells were washed, resuspended in FACS buffer and acquired on a Beckman Coulter CytoFlex S analyzer. The data was analyzed using FlowJo 10.

### Immune cell composition analysis by flow cytometry

All mice from the STING agonist-treated group (cGAMP-VLP and ADU-S100) and vehicle-treated group were sacrificed 24 hours after the last intratumoral injection. Spleen, draining/non-draining lymph nodes and tumors were excised. Splenocytes were isolated by pressing the spleen through a 40-µm cell strainer, axillary or inguinal LNs were dissected, pierced once with fine tip forceps, and collected into RPMI on ice. For the splenocytes, RPMI was replaced with 2 mL enzymatic solution of CO2-independent medium containing 1 mg/mL liberase (Sigma) and 20 μg/mL DnaseI (Roche), and incubated for 30 minutes in a 37°C incubator with gentle agitation. After 30 minutes, red blood cells were lysed using an ammonium chloride lysis buffer as described above. Cells were pelleted (300 x g, 10 minutes, 4°C) and resuspended in ice cold FACS buffer containing 0.5% BSA in PBS. Excised tumors were collected in RPMI supplemented with 10 % FCS and cut into small pieces. Tumor pieces were digested with 1 mg/mL liberase (Sigma) and 20 μg/mL DnaseI (Roche) with gentle continuous agitation (using mouse tumor dissociator gentleMACS). After 40 minutes digestion at 37°C, cells were passed through a 70-µm filter, washed by RPMI supplemented with 10 % FCS, and resuspended in FACS buffer. Single cells were stained according to standard protocols. Briefly, cells were surface-stained in 50 µL antibody-mix in FACS buffer: CD45.2 (clone 104), CD19 (clone 6D5), TCR-b (clone H57-597), CD4 (clone RM4-5), CD8 (Life Technologies), CD62L (clone MEL-14), CD69 (clone H1.2F3), CD44 (clone IM7), CD25 (clone PC61), NK1.1 (clone PK136), Nkp46 (clone 29A1.4), CD172a (clone P84), CD11b (M1/70), CD11c (Invitrogen), MHC-2 (clone M5/114.15.2), F4/80 (BM8), XCR1 (clone ZET), CD64 (clone X54-5/7.1), CD26 (clone H194-112) and CD86 (clone GL1). Dead cells were excluded using fixable viability stain according to the manufacturer’s instructions. For intracellular staining, cells were fixed for 30 minutes on ice using IC Fixation Buffer from Foxp3/Transcription Factor Staining Buffer Set, washed with 1X permeabilization buffer, stained and resuspended in FACS buffer containing ant-Foxp3 (clone FJK-16s) and anti-Ki67 (BD Biosciences). Single-cell suspensions were then analysed by flow cytometry using FACS LSRFortessa analyzer (BD Biosciences). For the analysis of the relative amounts of OST-STING in DCs and macrophages, splenocytes were stained with antibodies directed against CD11b (M1/70) and CD64 (clone X54-5/7.1), permeabilized with BD Cytofix/Cytoperm (BD Biosciences) for 30 min at 4°C, stained with 1/400 or 1/800 dilutions of Strep-Tactin APC (IBA GmbH) and analyzed by flow cytometry.

### LEGENDplex Assay

Serum samples were collected three hours after the first STING agonist injection and analyzed for inflammatory cytokines (IFN-α, IFN-ß, TNF-α and IL-6) using a LEGENDplex Mouse Inflammation Panel (BioLegend). For cell culture supernatants, IFN-α and IFN-β concentration were measured using a LEGENDplex Mouse Type 1/2 Interferon Panel (reference 740636). Data was acquired on a FACS Verse (BD Biosciences) and analyzed with BioLegend’s LEGENDplex Data Analysis Software. The standard curve regression was used to calculate the concentration of each target cytokine.

### Quantification and Statistical analysis

Statistical details of experiments are indicated in the figure legends, text or methods. Data were analyzed in GraphPad Prism 8 software. In Figures, * *P* < 0.05, ** *P* < 0.01, *** *P* < 0.001, **** *P* < 0.0001.

---

**Figure S1.**
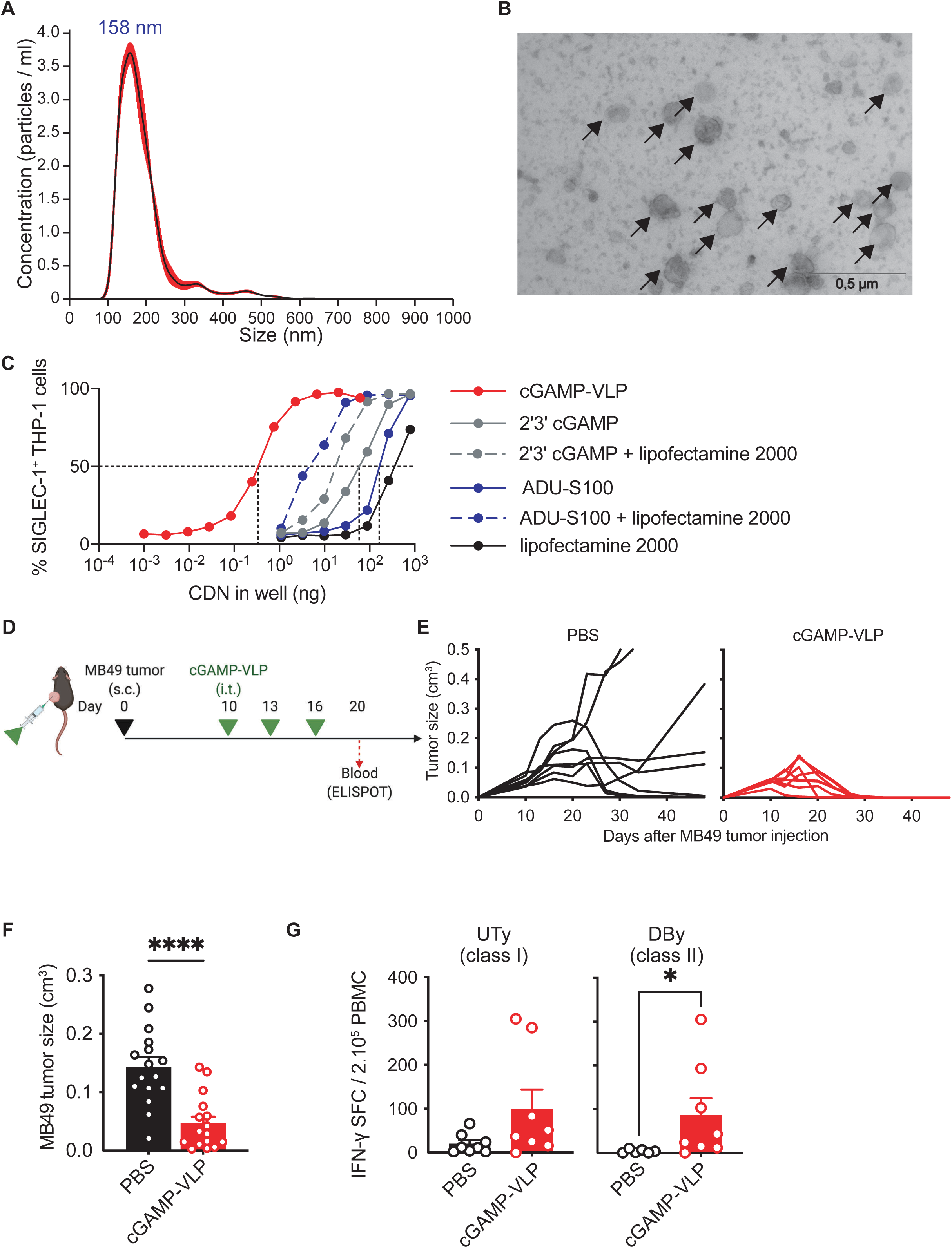
cGAMP-VLP induces antigen-specific anti-tumor immune responses by intra-tumoral injection. **(A)** Size distribution of purified cGAMP-VLP analyzed by Nanoparticle Tracking Analysis. Line at mean, red shading at 1 standard error of the mean (representative data of n = 21 experiments). **(B)** Electron microscopy image of purified cGAMP-VLPs. Scale bars at 0.5 µm. Arrows point to cGAMP-VLP. **(C)** SIGLEC-1 induction in THP-1 by increasing concentrations of cyclic dinucleotide (CDN) in the form of cGAMP-VLP, soluble 2’3’-cGAMP or soluble ADU-S100, with or without lipofectamine. Lipofectamine 2000 alone condition is plotted at the doses equivalent to the conditions with CDN. Dotted lines indicate CDN dose at 50% SIGLEC-1^+^ cells. **(D)** Overview of the experimental design. Treatments were started on tumors of 50 mm^3^ average volume per group at day 10. Mice were treated at days 10, 13 and 16 with cGAMP-VLP or PBS injected by the i.t. route. **(E)** Growth curves of individual MB49 tumors (n = 8 mice per group). **(F)** Size of tumor 17 days after tumor implantation in treated mice (line at mean + SEM, n = 12 mice per group combined from 2 independent experiments, Mann-Whitney test). **(G)** T cell responses against UTy (class I peptide) and DBy (class II peptide) in blood of mice 20 days after tumor implantation, assess by IFN-γ ELISPOT (bar at mean + SEM, n = 6 to 8 mice per group, Mann-Whitney test).

**Figure S2.**
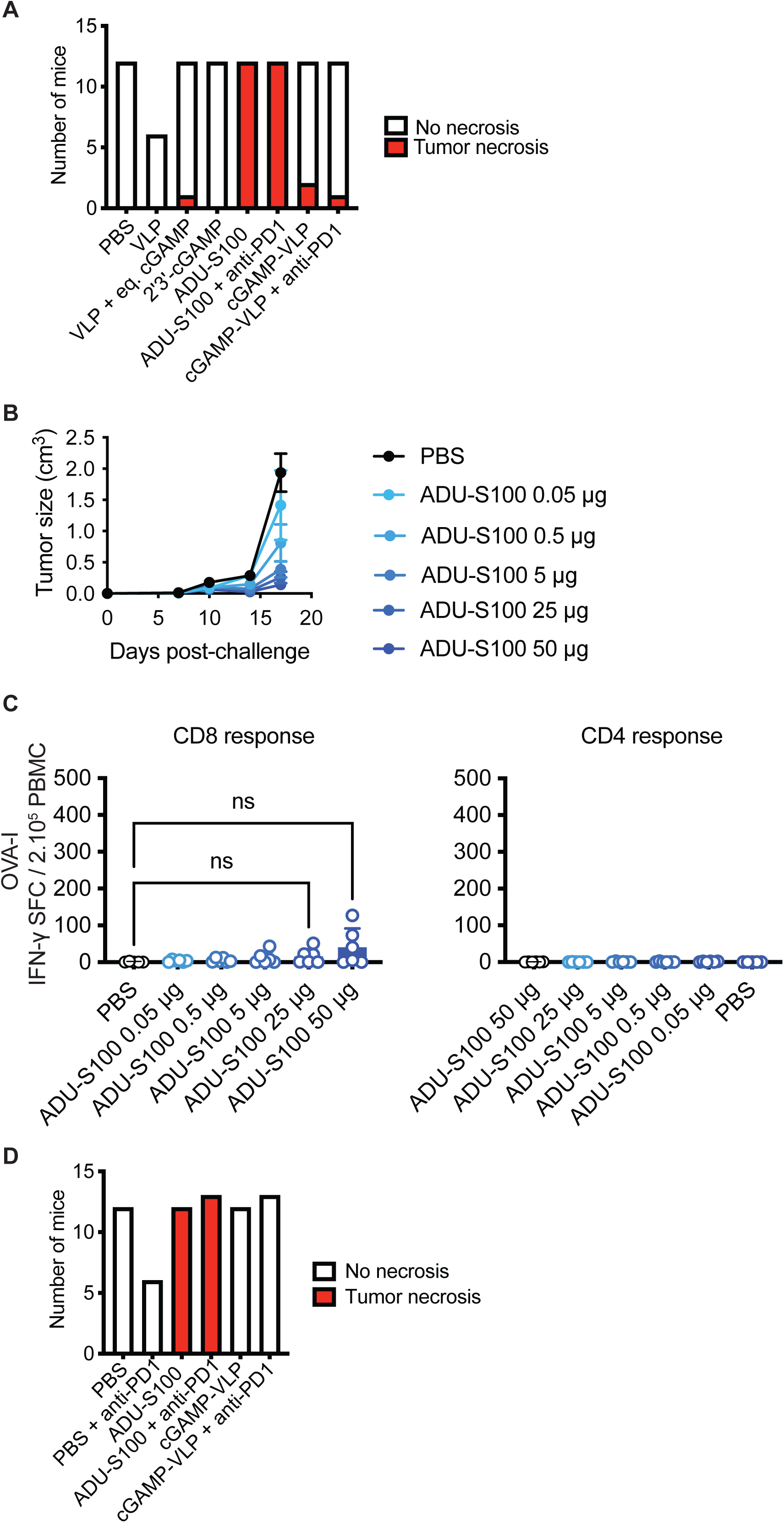
Responses to lower doses of ADU-S100 and tumor necrosis. **(A)** Number of tumor necrosis events after the indicated treatments in single tumor experiments. **(B)** Mean growth over time of B16-OVA tumors treated as indicated by different doses of ADU-S100 (line at mean + SEM, n = 5 to 6 mice per group). **(C)** Ova-specific CD8 (OVA-I) and CD4 (OVA-II) T cell responses in blood, assess by IFN-γ ELISPOT (bar at mean + SEM, n = 5 to 6 mice per group, Kruskal-Wallis test with Dunn post-test). **(D)** Number of necrosis events in the injection tumor after the indicated treatments in dual tumor experiments.

**Figure S3.**
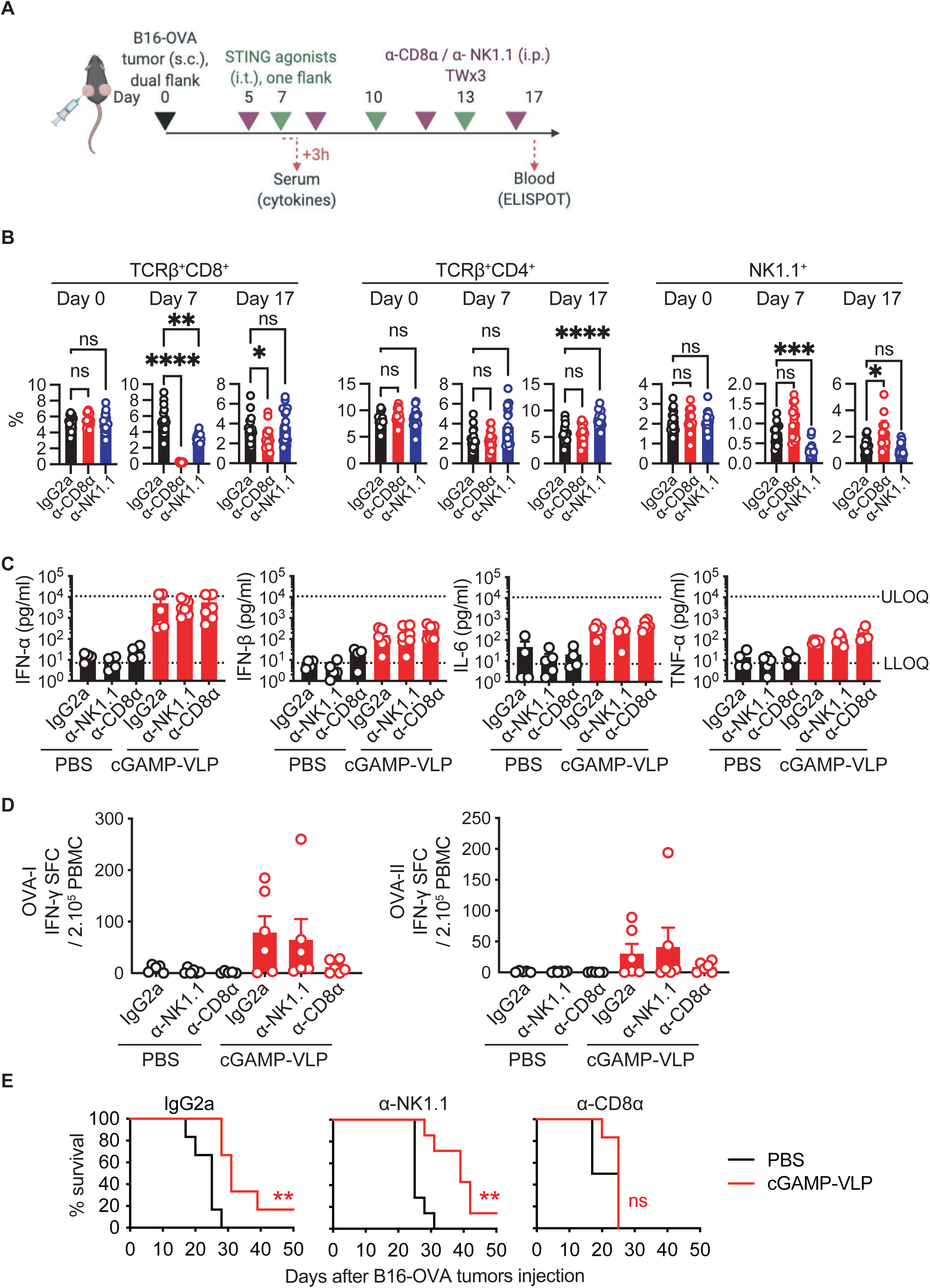
The anti-tumor effect of cGAMP-VLP requires CD8^+^ T lymphocytes but not NK cells. **(A)** Evaluation of the role of CD8^+^ T cells and NK cells, overview of the experiment. Mice were randomized at day 7 and treated by i.t. injection at days 7, 10 and 13 with PBS, 50 µg ADU-S100 or 50 ng cGAMP-VLP. Treatments were initiated on palpable tumors **(B)** Fraction of CD8^+^ T cells, CD4^+^ T cells and NK cells in the blood at days 0, 7 and 17 after injection with isotype, anti-CD8βα or anti-NK1.1 (bar at mean + SEM, n = 18 to 21 mice per group, Kruskal-Wallis with Dunn post-test). **(C)** Concentrations of IFN-α, IFN-β, IL-6 and TNF-α in the serum of B16-OVA dual tumor-bearing mice 3 hours after first injection of cGAMP-VLP or PBS, in mice treated with antibodies as indicated (bar at mean + SEM, n = 4 to 7 mice per group, LLOQ = lower limit of quantification, ULOQ = upper limit of quantification). **(D)** Ova-specific CD8 (OVA-I) and CD4 (OVA-II) T cell responses in blood of mice treated as indicated, 17 days after tumor implantation, assess by IFN-γ ELISPOT (bar at mean + SEM, n = 5 to 6 mice per group). **(E)** Survival of B16-OVA dual tumor-bearing mice treated as indicated (log-rank Mantel-Cox test).

**Figure S4.**
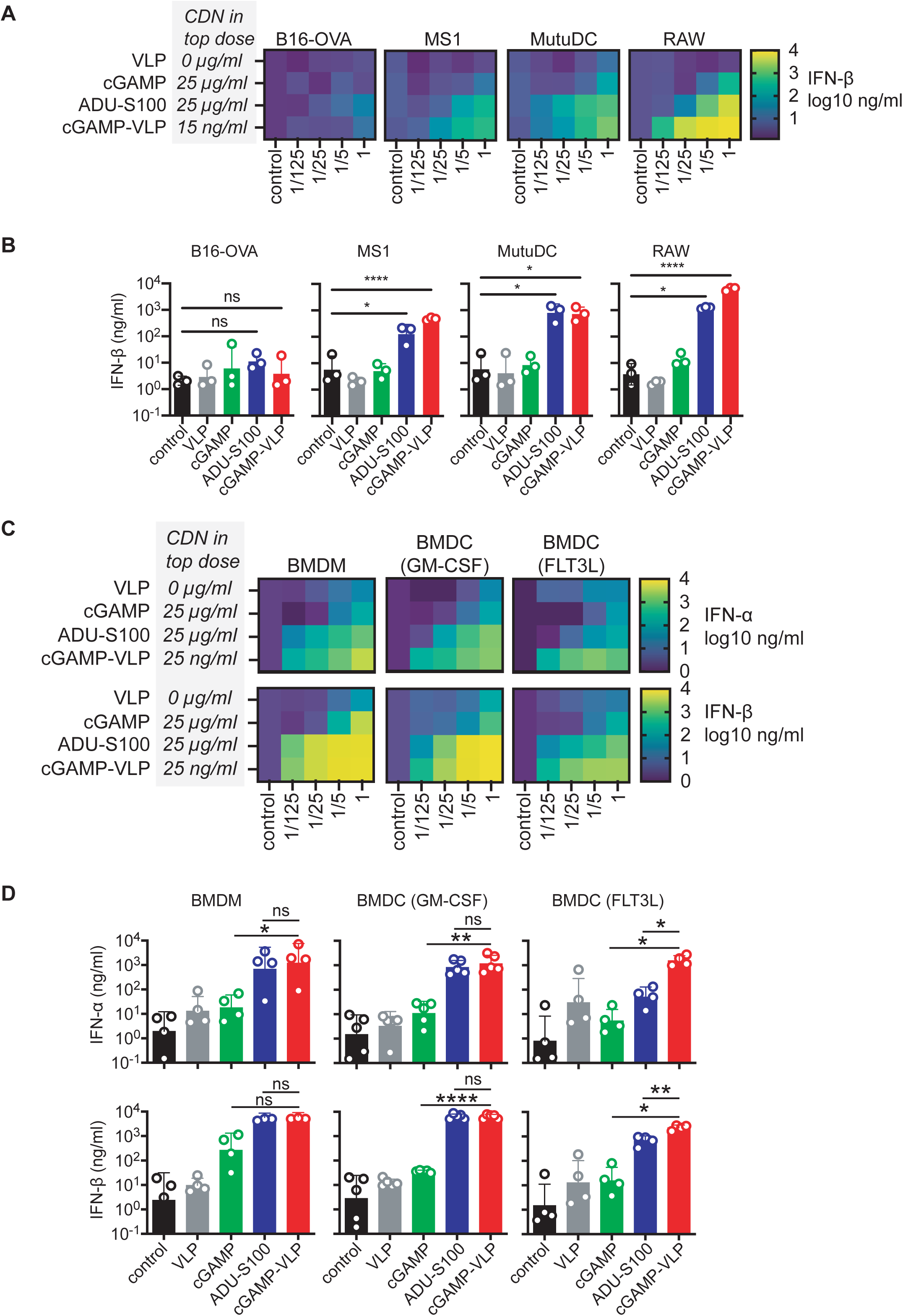
Response of cell lines and dendritic cells to cGAMP-VLP. **(A)** Production of IFN-β by B16-OVA, MS1, MutuDC and RAW cell lines after stimulation with dose titration of VLPs, cGAMP, ADU-S100 and cGAMP-VLP starting at the indicated top dose (averages from n=3 independent experiments). **(B)** Statistical analysis of IFN-β at dilution 1/5 (bar at mean + SEM, n=3 independent experiments, one-way ANOVA with Tukey post-test on log-transformed data). **(C)** Production of IFN-α and IFN-β by BMDM, BMDC (GM-CSF) and BMDC (FLT3L) after stimulation with dose titration of VLPs, cGAMP, ADU-S100 and cGAMP-VLP at the indicated top dose (averages from n=4 or 5 independent experiments). **(D)** Statistical analysis of IFN-α and IFN-β at dilution 1/5 (bar at mean + SEM, n=4 or 5 independent experiments, one-way ANOVA with Tukey post-test on log-transformed data).

**Figure S5.**
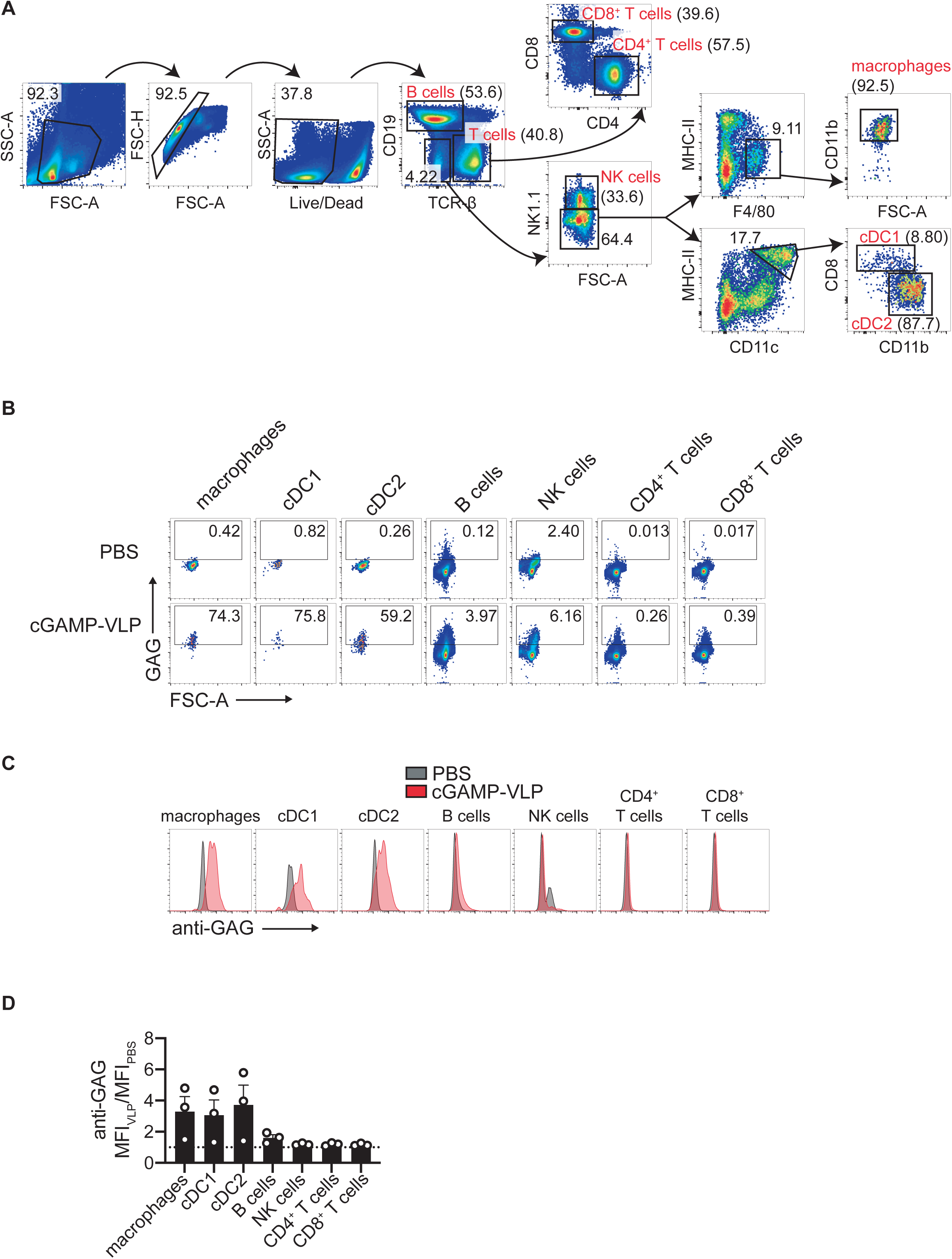
Capture of cGAMP-VLP by splenocytess. **(A)** Gating strategy of immune cells subsets for cGAMP-VLP capture experiments (representative of n=3 independent experiments). **(B)** Anti-GAG staining and forward scatter in the indicated immune cells from splenocytes treated with PBS or cGAMP-VLP (representative of n=3 independent experiments). **(C)** Overlaid anti-GAG staining in the indicated immune cells from splenocytes treated with PBS or cGAMP-VLP (representative of n=3 independent experiments). **(D)** Ratio of anti-GAG mean fluorescence intensity for cGAMP-VLP over PBS (bar at mean + SEM, n=3 independent experiments).

**Figure S6.**
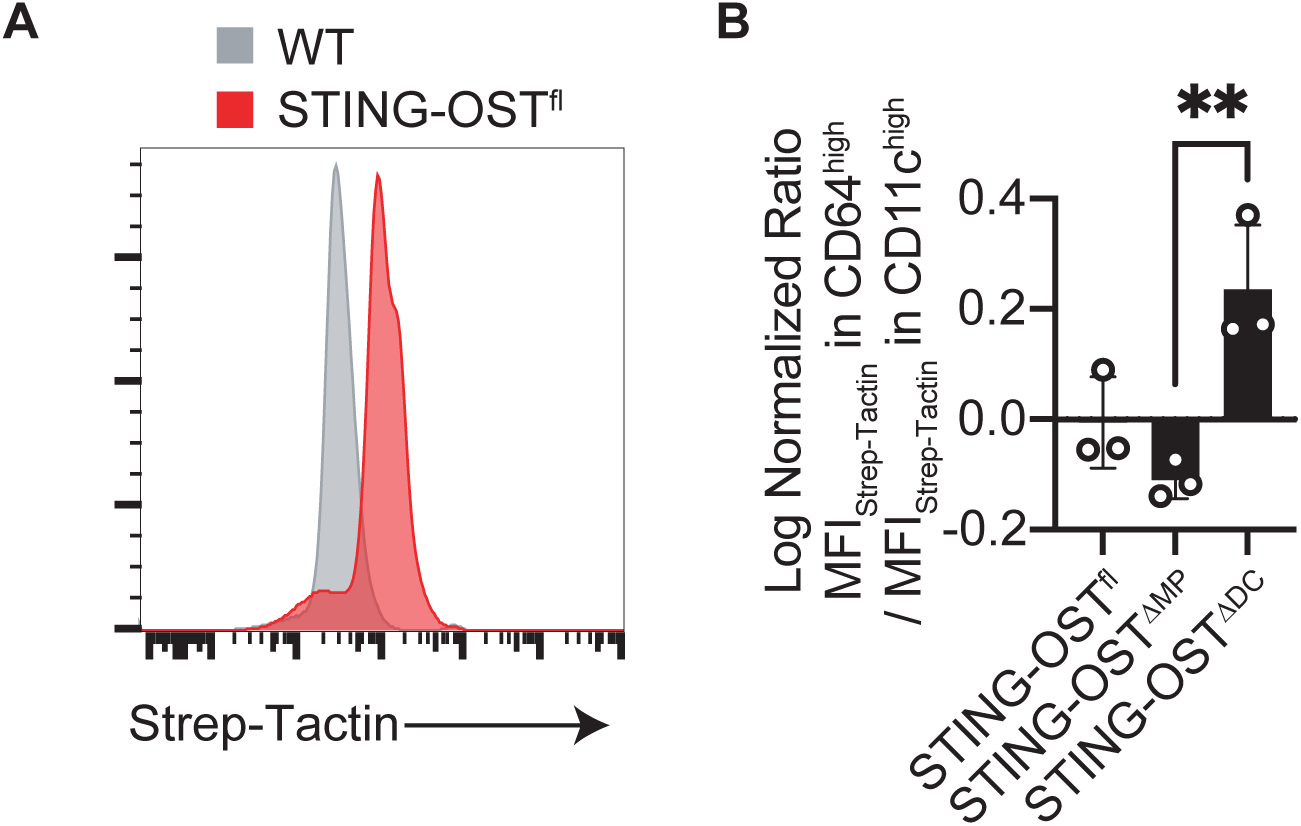
Preferential deletion of STING in macrophages or dendritic cells. **(A)** Representative Strep-Tactin staining in total live single cells in spleen of WT and STING-OST^fl^ mice. **(B)** Relative Strep-Tactin staining in CD64^high^ and CD11c^high^ live single cells in spleen of the indicated mouse strains (n=3 combined from 2 independent experiments, ANOVA with Tukey test).

**Figure S7.**
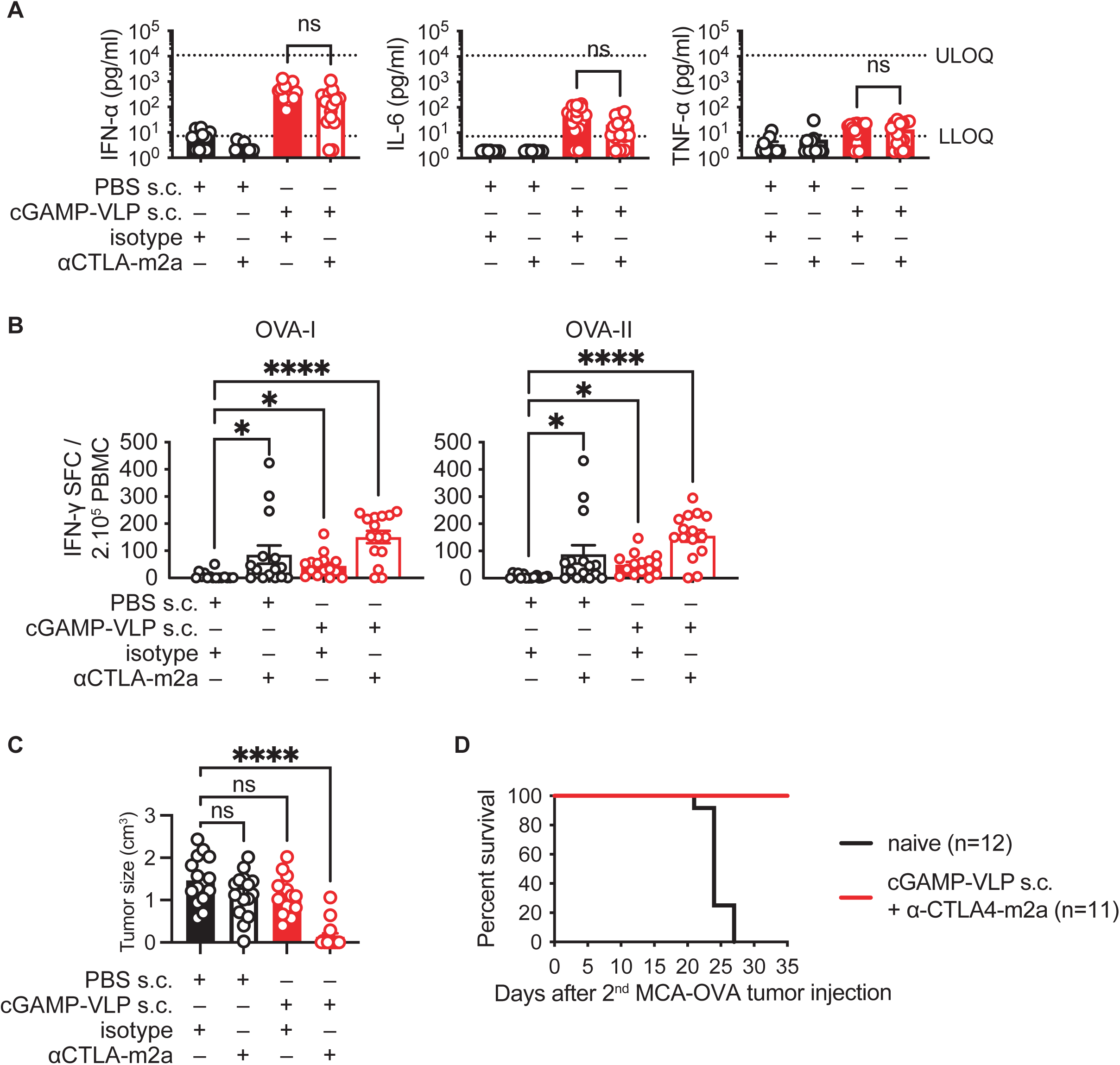
Additional results for the response to cGAMP-VLP combined with anti-CTLA-m2a. **(A)** Concentrations of IFN-α, IL-6 and TNF-α in the serum of MCA-OVA tumor-bearing mice 3 hours after the first treatment with PBS or 50 ng cGAMP-VLP injected by the s.c., and i.p. injection of αCTLA4-m2a or isotype. Treatments were started on tumors of 50 mm^3^ average volume per group (bar at mean + SEM, n = 11 to 15 mice per group, combined from 2 independent experiments, Kruskal-Wallis with Dunn post-test, LLOQ = lower limit of quantification, ULOQ = upper limit of quantification). **(B)** Ova-specific CD8 (OVA-I) and CD4 (OVA-II) T cell responses in blood of mice 16 days after tumor implantation, assess by IFN-γ ELISPOT (bar at mean + SEM, n = 15 mice per group, combined from 2 independent experiments, Kruskal-Wallis with Dunn post-test). **(C)** Size of tumor 28 days after tumor implantation in treated mice (line at mean + SEM, n = 15 mice per group combined from 2 independent experiments, Mann-Whitney test). **(D)** Survival of mice after secondary challenge. In complete responding mice, MCA-OVA cells were injected 55 days from the first injection of tumor cells and treatments (combined from 2 experiments, Mantel-Cox test).

